# Snapshot of in-cell protein contact sites reveals new host factors and hijacking of paraspeckles during influenza A virus infection

**DOI:** 10.1101/2025.03.09.642134

**Authors:** Iuliia Kotova, Lars Mühlberg, Konstantin Gilep, Dingquan Yu, Daniel Ziemianowicz, Stephanie Stanelle-Bertram, Sebastian Beck, Kyungmin Baeg, Olivier Duss, Gülsah Gabriel, Fan Liu, Boris Bogdanow, Jan Kosinski

**Affiliations:** European Molecular Biology Laboratory Hamburg, Notkestraße 85, 22607 Hamburg, Germany; Centre for Structural Systems Biology (CSSB), Notkestraße 85, 22607 Hamburg, Germany; Department of Structural Biology, Leibniz-Forschungsinstitut für Molekulare Pharmakologie (FMP), Berlin, Germany; Research Department Viral Zoonoses – One Health, Leibniz Institute of Virology (LIV), Martinistraße 52, 20251 Hamburg, Germany; Molecular Systems Biology Unit, European Molecular Biology Laboratory, Meyerhofstrasse 1, Heidelberg 69117, Germany; Research Center for Emerging Infections and Zoonoses, University of Veterinary Medicine Hannover, Buenteweg 17, 30559 Hannover, Germany; Institute of Virology, Charité—Universitätsmedizin Berlin, Freie Universität Berlin and Humboldt-Universität zu Berlin, Berlin Institute of Health, Berlin, Germany

**Author notes:** Correspondence (J.K), (B. B.), (F.L).

**Keywords:** Influenza virus, protein-protein interactions, host-pathogen interactions, cross-linking mass spectrometry, AlphaFold, host factors, membrane, phase-separation, paraspeckles, NEAT1, hemagglutinin, LAT1

## Abstract

Influenza A virus (IAV) hijacks host cellular machinery, but many virus–IAV interactions and contacting protein sites remain uncharacterised, particularly those dependent on intact cellular architecture, such as membrane-associated or phase-separated compartments. Here, we applied in-cell cross-linking mass spectrometry (XL-MS), integrated with AlphaFold-based structural modelling and functional assays, to map protein-protein contact sites in IAV-infected human cells. This approach revealed previously unrecognised virus–host interactions linked to spatially organised processes, including the maturation pathway of HA through the membrane-bound ER– Golgi system, the novel interaction of M2 with the membrane-embedded LAT1 amino acid transporter, and the progressive disassembly of paraspeckles–phase-separated compartments in the nucleus. We validate M2-LAT1 interaction and paraspeckle disassembly in human primary lung epithelial cells and show that the paraspeckle disassembly constitutes a new and unique infection mechanism through which IAV releases RNA-binding proteins that support viral RNA replication. These findings advance the understanding of IAV manipulation of host cellular processes and illustrate how the integrative in-cell structural system biology approach captures native host-pathogen interactomes, infection pathways, and host cell perturbations.

## Main

Influenza A virus (IAV) remains a major global health threat, causing substantial annual morbidity and mortality^1^. The viral replication cycle culminates in the biogenesis of viral progeny within infected host cells, driven by protein-protein interactions (PPIs) between viral and host proteins^2^. The IAV genome consists of eight gene segments encoding up to 14 proteins, depending on the strain^2^. These proteins engage in dynamic interactions that form and dissociate depending on the stage of infection and are often confined to specific cellular compartments and organelles. Thus, understanding host-IAV PPIs in their native cellular and spatial context is essential for elucidating viral strategies of host manipulation and identifying therapeutic targets.

One of the most prominent examples of the importance of subcellular context in defining IAV-host PPIs is viral RNA (vRNA) synthesis. This process is confined to the nucleus and is catalysed by the viral RNA-dependent RNA polymerase complex (RdRp), composed of PA, PB1, and PB2 proteins. vRNA synthesis is further embedded in the spatial context of the viral ribonucleoproteins (vRNPs), composed of RdRp and viral vRNA wrapped around NP protein filament^2^. Moreover, assembly of RdRp, along with transcription and replication, is facilitated by the recruitment of additional host factors^3–6^.

Similarly, nuclear vRNA processing is modulated by context-specific PPIs between viral proteins, such as NS1 and NP, and host RNA-binding proteins^7,8^. These proteins remodel sub-nuclear compartments, such as nuclear speckles, which are essential for splicing of IAV mRNA^9^. Furthermore, the viral PA-X nuclease, expressed from an alternative reading frame of PA^10^, degrades nuclear host transcripts^11,12^, which are known to organise subnuclear compartments in uninfected cells^13^. However, how context-specific protein and RNA interactions in infected cells influence subnuclear compartments and whether this impacts replication remains poorly understood.

The importance of context-specific PPIs within dynamic macromolecular assemblies and organelles extends beyond the nucleus. Newly synthetised vRNPs are exported from the nucleus by nuclear export factors, a process mediated by viral proteins M1^14^ and NEP^15^. Consequently, vRNPs are trafficked through the cytoplasm to the plasma membrane via an interaction with host RAB11A protein^16^ at the endoplasmic reticulum (ER), which undergoes architectural reorganisation during infection^17^. In parallel, surface viral glycoproteins hemagglutinin (HA) and neuraminidase (NA) are synthesised in the ER, glycosylated in the Golgi, and trafficked to the plasma membrane^18^, involving extensive interactions with the cellular machinery for glycoprotein maturation. Ultimately, glycoproteins, vRNPs, M1 and the ion channel M2 converge to assemble into viral particles that bud from the plasma membrane.

Despite the clear importance of subcellular and infection contexts in shaping functionally relevant PPIs during IAV infection, systematic data on IAV-host PPIs in these native contexts is lacking. Instead, methods such as affinity purification mass spectrometry (AP-MS) and yeast-two-hybrid (Y2H) assays have reported many candidate IAV-host PPIs^19–28^, but have relied on cell lysates disrupting native cellular architecture and dynamic macromolecular assemblies and often used non-infected cells. Moreover, these methods do not provide insights into interaction sites and structures of complexes, which have barely been characterised at atomic resolution for IAV host-pathogen PPIs.

Here, we applied in-cell cross-linking mass spectrometry (XL-MS)^29^ combined with AlphaFold-based structural modelling and functional analysis to map IAV-host PPIs in infected human lung epithelial cells. In-cell XL-MS captures interactions directly in intact cells and defines residue-to-residue contact sites between proteins in close proximity (within 40 Å, based on the distance constraint of disuccinimidyl sulfoxide (DSSO) cross-linker used). This approach identified hundreds of cross-links between IAV and human proteins in their native context.

Integrating XL-MS with structural modelling and functional screens, we uncovered previously unrecognised virus-host interactions and host factors linked to spatially organised processes. These findings include compartment-specific contacts along the HA maturation pathway in the ER/Golgi system, the identification of the LAT1 amino acid transporter as a membrane-associated interactor of M2, and the disruption of paraspeckles—membraneless nuclear organelles—through extensive interactions with host RNA-binding proteins. The disruption of paraspeckles has not been previously demonstrated for IAV or any other virus. Mechanistically, we propose that NP and NS1 interactions with paraspeckle protein components, degradation of non-coding RNA NEAT1_2 by PA-X, and RNA Pol II inhibition collectively drive this process. This disruption liberates RNA-binding proteins that facilitate vRNA synthesis and replication.

Together, our findings provide a spatially-resolved snapshot of IAV-host contact sites in intact cells, integrating structural data and functional analysis to reveal new mechanisms by which the virus reprogrammes host cellular machinery.

## Results

### Mapping IAV-human contact sites using in-cell cross-linking

To map PPIs and their contact sites in IAV-infected cells, we applied XL-MS-based structural host-virus interactome profiling (SHVIP) (Fig. 1a). SHVIP mitigates the inherent sensitivity limitation of conventional in-cell XL-MS by combining XL-MS and bio-orthogonal labelling with unnatural amino acids for the selective enrichment of newly synthesised viral proteins^29^.

**Fig. 1.**
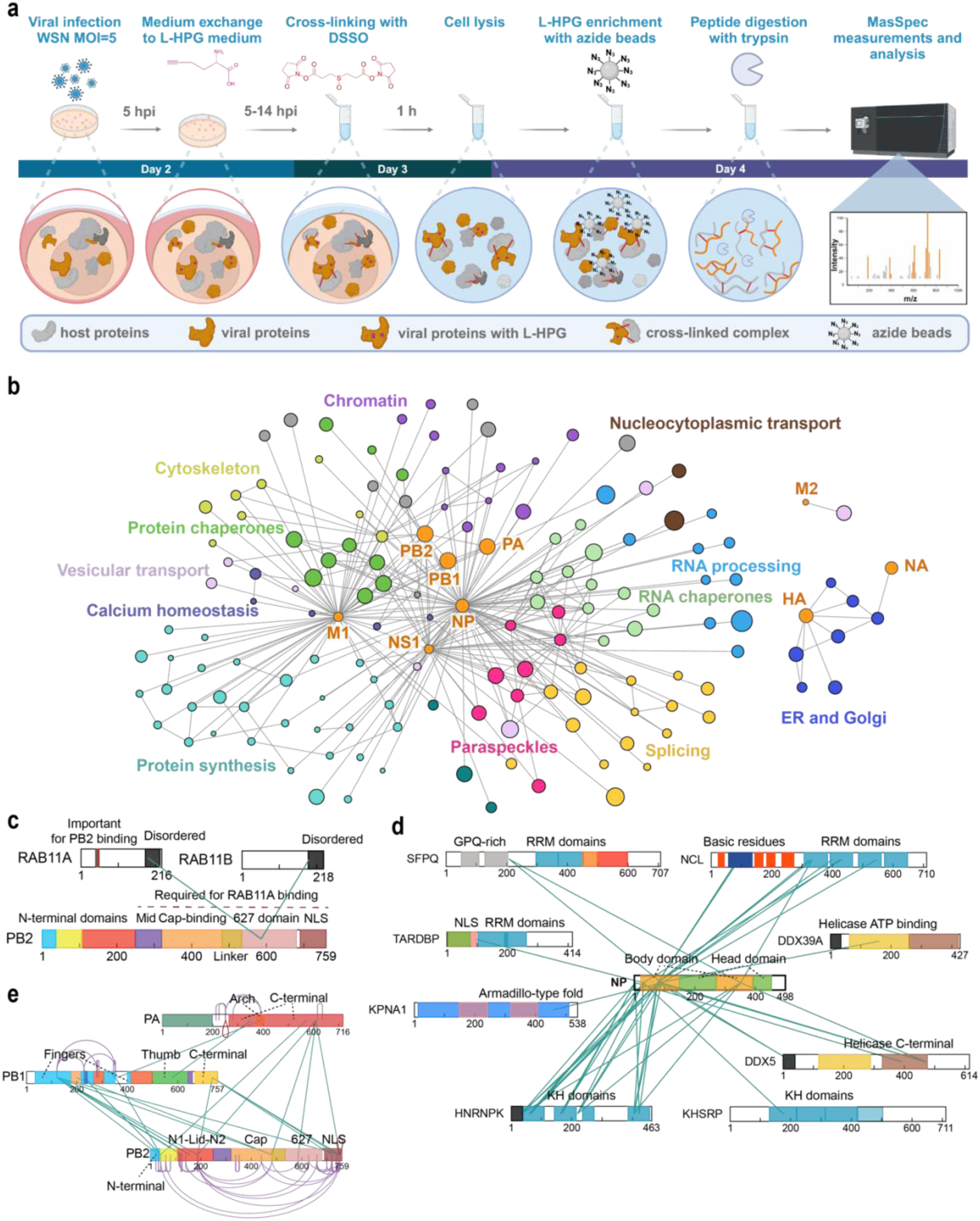
In-cell cross-linking during influenza A virus infection. **a,** Schematic of in-cell cross-linking using SHVIP. IAV-infected cells are labelled with HPG from 5 to 14 hpi, followed by DSSO cross-linking. After quenching, L-HPG-incorporated proteins are enriched via click chemistry, digested with trypsin, and cross-links are analysed by MS. *Created in BioRender.* https://BioRender.com/g57m779. **b**, Virus-centric cross-linking network at a 2% FDR threshold. Viral proteins are shown in orange, and host proteins are coloured by their functional category. Only host proteins linked to viral proteins are shown with names omitted for clarity. The network including host names is shown in Extended Data Fig. 2c, Interaction between PB2 and RAB11A/B, consistent with the established binding domain of PB2. Interlinks are shown in green, while intralinks are omitted for clarity. **d**, Cross-links between NP and established host factors involved in influenza virus infection. Colour shading represents distinct structural domains, with names shown only for those forming cross-links for clarity. Interlinks are shown in green, while intralinks are omitted for clarity. **e**, Cross-links between subunits of vRdRp. Interlinks are shown in green, while Intralinks in violet.

To apply SHVIP to IAV, we optimised the labelling to enrich the viral proteome while preserving the physiological context of infected cells (Extended Data Fig. 1a-e). Previous bio-orthogonal amino acid labelling studies showed that dominant IAV protein synthesis begins a few hours post-infection and slows during later stages^30^.

We timed the cross-linking and labelling scheme to capture the late infection stage (see Methods and Extended Data Fig. 1c-f), during which vRNP export, cytoplasmic genome transport, and virion assembly occur at the plasma membrane^31^. Additionally, we confirmed that L-HPG labelling did not affect cell viability or infection progression (Extended Data Fig. 1g-i), ensuring minimal disruption to productive IAV infection.

To confirm that SHVIP efficiently enriches the viral proteome, we compared host and viral protein abundance in enriched samples versus input aliquots across three replicates. Viral proteins were, on average, ten times more abundant in enriched samples (Extended Data Fig. 1i-l). Likewise, proteomic XL-MS analysis showed that viral protein pairs and cross-links were detected 2-10 times more frequently in enriched samples (Extended Data Fig. 1m-o). Pairwise reproducibility across replicates (47% on average) (Extended Data Fig. 1p-r) matched previous large-scale XL-MS studies^32^ and the SHVIP HSV-1 dataset^29^.

### Extensive *in-cell* cross-linking network

We analysed the combined raw data from three biological SHVIP replicates, applying a consecutive 2% FDR threshold for cross-link spectrum match (CSM), residue-pair (Extended Data Table 1a) and PPI levels (Extended Data Table 1b). This dataset included 13201 unique residue-pair cross-links (Extended Data Table 1c) originating from 1961 proteins. In particular, 111 viral-viral cross-links and 877 viral-host cross-links were identified. These involve nine viral proteins, 141 host proteins, and 200 viral-host protein pairs (Extended Data Table 1d, Fig. 1b).

Of the viral-host interactions, at least 50 protein pairs involved host proteins known to bind viral proteins (Extended Data Table 2, Extended Data Fig. 2). We also found cross-links to host factors known to function in the IAV infection cycle, such as RAB11A (Fig. 1c), KHSRP^33^ and TDP-43(TARDBP)^34^ (Fig. 1d), ANP32A and ANP32B^35^, nucleosomes^36,37^, microtubules^38^, and chaperonins^39^.

The captured interactome is enriched for functional Gene Ontology (GO) terms related to gene expression, RNA processing, splicing, and transport, response to unfolded proteins, protein synthesis, ribosome biogenesis, chromatin organisation, and regulation of calcium homeostasis and phosphorylation (Fig. 1b, Extended Data Fig. 2, Extended Data Table 3). The interacting proteins localise to both nuclear and cytoplasmic compartments, with notable enrichment in the nucleolus, cytoskeleton, and vesicles (Extended Data Table 3). Many of these enriched spatial and functional categories align with previous influenza virus PPI studies^19,40^.

To demonstrate the benefit of our approach over previous IAV PPI studies, we compared subcellular localisation of viral proteins (Extended Data Fig. 3a), cross-linked host proteins (Extended Data Fig. 3b), and interactors reported by Watanabe & Kawakami et al. (2014)^19^ (Extended Data Fig. 3c), Haas et al. (2023)^21^ (Extended Data Fig. 3d), and the meta-analysis by Chua et al. (2022)^40^ (Extended Data Fig. 3e). Strikingly, SHVIP shows a more specific and biologically consistent localisation pattern, with cross-linked host proteins localising to compartments matching those of the respective viral proteins. In contrast, the earlier PPI datasets frequently report localisations incompatible with the known viral protein localisations—for example, widespread mitochondrial interactors across nearly all viral proteins, despite no clear evidence for mitochondrial localisation of IAV proteins other than PB2^41,42^. Similarly, the other studies erroneously reported that ER- and plasma-membrane-associated proteins such as M2, HA, and NA interact with numerous nuclear proteins. These discrepancies likely reflect artefacts introduced by overexpression and biochemical purification, which can promote non-physiological interactions. In contrast, SHVIP preserves native infection context and subcellular architecture, and thus yields a more reliable and spatially resolved interaction landscape.

Another unique advantage of the XL-MS over AP-MS and Y2H is that cross-links provide distance constraints to corroborate protein structures or guide structural modelling^43^. Mapping host-host cross-links onto structures from the Protein Data Bank (PDB) showed that 93% of the Cα-Cα distances between cross-linked residues fall within the DSSO cross-linker constraint of 40 Å (Extended Data Fig. 4). For viral complexes such as RdRp, cross-link satisfaction rates are difficult to calculate due to conformational and oligomeric heterogeneity^44,45^. Nonetheless, the cross-linking pattern of RdRp (Fig. 1e) aligns with the cross-linking pattern observed for purified RdRp^45^. For IAV-host interactions, no structures available in the PDB contained cross-links that connected the structurally resolved regions of IAV and human proteins, underscoring the challenges of capturing these interactions structurally. Several IAV-human interactions were modelled with good confidence using AlphaFold2-multimer^46,47^, AlphaFold 3^48^ and the cross-link-guided modelling protocol AF3x^49^ (Extended Data Fig. 6, Extended Data Table 4). In contrast, many other pairs could not be confidently modelled—likely explained by the reliance of AlphaFold on co-evolutionary signals and sequence pairing from the same species, which is not achievable in host-pathogen systems^47^. The successful models indicate direct physical interactions, identify candidate contact residues, and further corroborate that our map represents a snapshot of IAV–human residue contact sites in intact cells.

Overall, the substantial overlap of the identified cross-links with functional host protein categories, known host factors, PPIs characteristic of IAV infection, and AlphaFold models—together with the consistent subcellular localisation of the interacting host proteins—demonstrates that SHVIP delivers a structurally informative IAV-host interactome, and, in contrast to previous PPI studies, it captures interactions in their native cellular context.

### Cross-linking network and siRNA screening reveal new host factors and functional hypotheses

To demonstrate that our cross-linking network identifies novel host factors in IAV infection and generates a wealth of functional hypotheses, we performed an siRNA-mediated loss-of-function screen targeting selected cross-linked host proteins (Fig. 2a). We prioritised host proteins with at least three cross-links to viral proteins, excluding well-studied host factors. Additionally, we included six host proteins with fewer cross-links—ERP29, LMAN2, EEF1A1, SET, SSB, and RALY—due to their roles in nucleic acid metabolism, gene expression regulation, and ER localisation. We also included SLC3A2 as the sole identified M2 interactor, and RAB11A (a known IAV dependency factor) as a positive control. IAV infection efficiency in siRNA-treated cells was quantified using a luciferase assay^50^ (Fig. 2a, Extended data Fig. 5a-b). RAB11A reduced viral replication by ~10-fold (Fig. 2b), which is consistent with previous reports^31,51^ and validates our assay. In addition, knockdown of ribosomal 60S subunit proteins RPL35 (cross-linked to NS1, M1, and NP) and RPL10A (cross-linked to M1) significantly reduced viral replication, whereas knockdown of other ribosomal proteins had no effect on viral titres, suggesting that these interactions have a specific function (Fig. 2b, Extended data Fig. 5a-b). These findings are consistent with the role of NS1 in regulating protein translation^52^ and suggest additional roles for IAV interactions with the translational machinery.

**Fig. 2.**
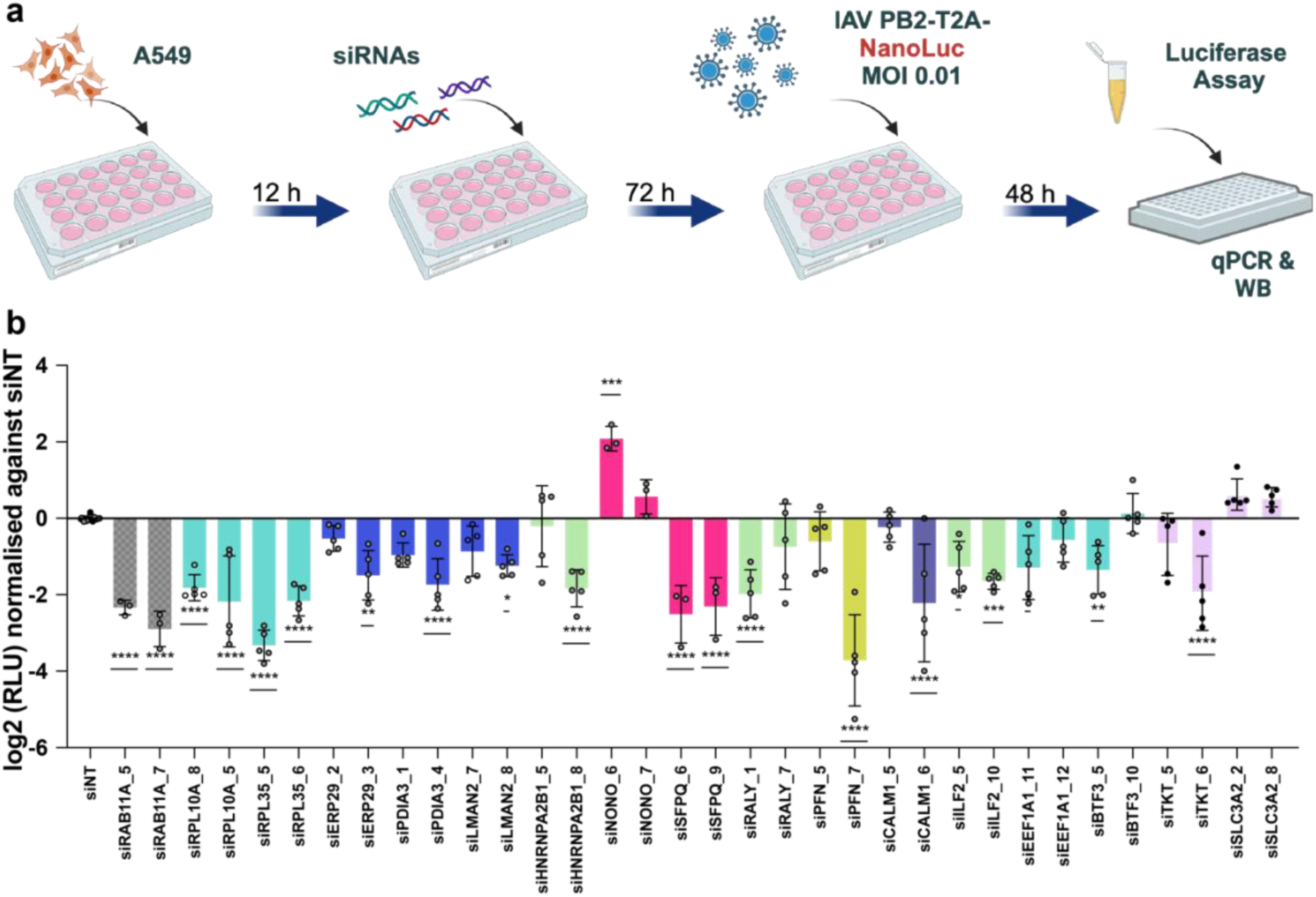
Impact of siRNA-mediated knockdown on IAV infection. **a**, Schematic of luciferase assay: A549 cells were transfected with siRNAs and at 72 hpt (hours post-transfection), cells were infected with recombinant WSN virus expressing PB2-T2A-NanoLuc. After 48 hours, luciferase activity was measured (Extended data Fig. 5C-D), providing read-out proportional to viral polymerase levels in infected, siRNA-treated cells^52^. Created in BioRender https://BioRender.com/h91s731. **b,** Log2-transformed relative luciferase units [Log2 (RLU)] normalised to a non-targeting siRNA control (siNT). Data are represented as mean ± standard deviation (SD) (n = 3 to 5 Statistical significances: *p<0.05, **p<0.01, ***p<0.001, ****p<0.0001 (one-way ANOVA and Dunnett’s multiple comparisons test, reference: siNT). All statistical tests were two-sided unless otherwise specified. See also Extended data Fig. 5A-B.

We further examined SLC3A2, the knockdown of which caused a small increase but was statistically significant only for one of the two siRNAs (Fig. 2b). SLC3A2, as well as its known interactor SLC7A5 (both are components of a dimeric complex, namely the large neutral amino acid transporter (LAT1)^53^ cross-linked to M2 with two cross-links at 2% FDR and an additional cross-link at 5% FDR (Extended Data Table 1e-g). All cross-links are consistent with the orientation of LAT1 and M2 across the membrane (Fig. 3a). LAT1 is essential for amino acid uptake and mTORC1 activation, processes supporting cell growth and survival^53,54^. Knockdown of either LAT1 subunit increased IAV infection in A549 cells (Fig. 3b). To validate the interaction, we performed co-IPs from infected cells against SLC3A2 and SLC7A5. M2 co-immunoprecipitated with both LAT1 subunits (Extended Data Fig. 7a). Additionally, we performed an AP-MS experiment in infected cells comparing pull-down against SLC7A5 and M2 to isotype-matched specificity controls. This revealed the mutual enrichment of M2 and SLC7A5 (Fig. 3c, Extended Data Fig. 7B). Co-localisation of M2 and LAT1 at the plasma membrane was observed in A549 (Fig. 3d-f) and primary human bronchial epithelial cells (HBEpCs) (Extended Data Fig. 7d-h), indicating that the interaction between M2 and LAT1 subunits occurs at cellular membranes and is preserved in the physiological context of primary cells. These findings suggest potential roles of this interaction in regulating amino acid transport or M2 localisation.

**Fig. 3.**
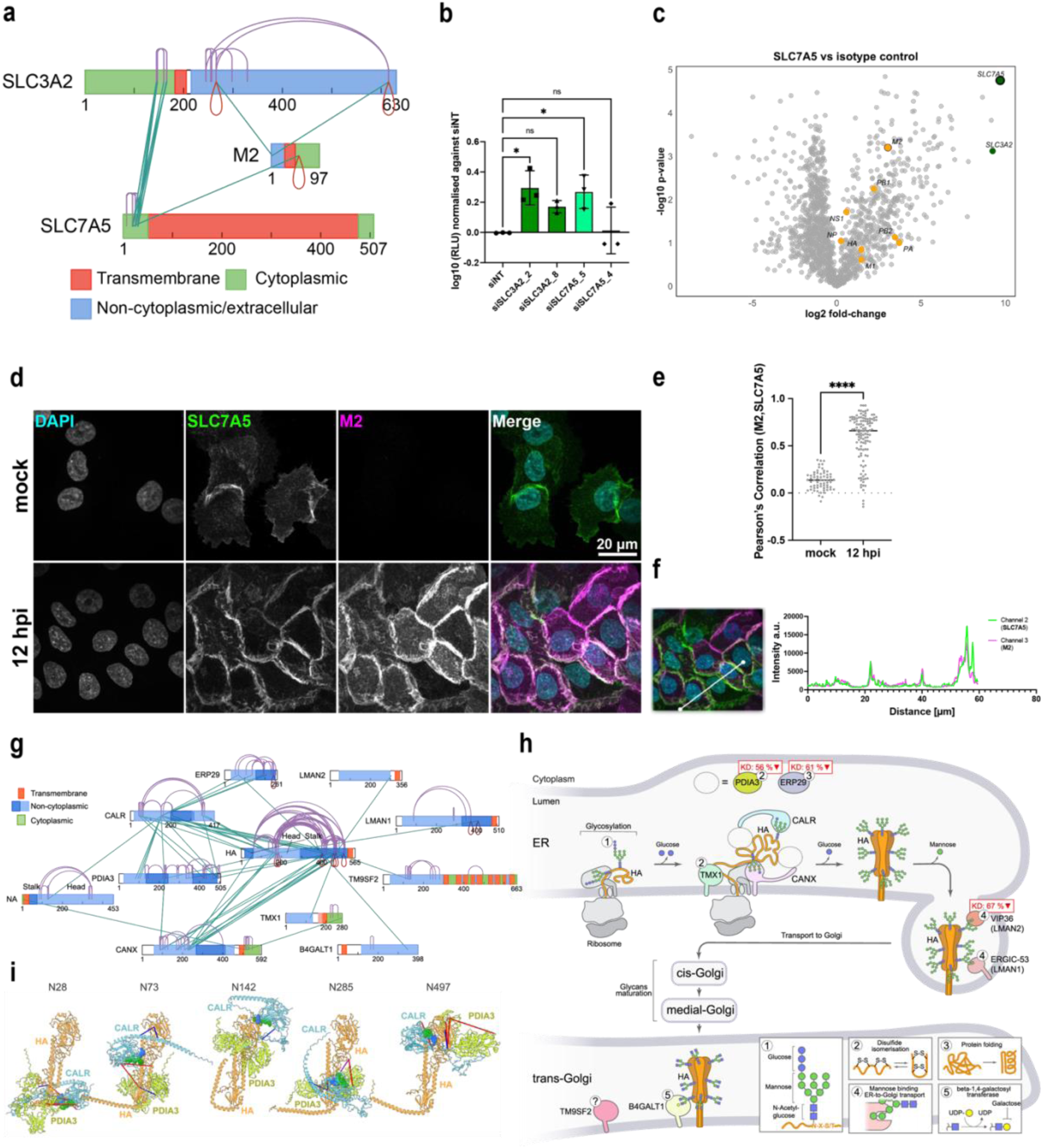
Analysis of interactions at membrane compartments. **a,** Cross-linking network of M2 and LAT1. **b**, Impact of LAT1 knock-down on infection presented as Log(RLU) normalised to siNT. Data are represented as mean ± SD (n = 3), with individual experiment means shown as circles. Statistical analysis was performed using ordinary one-way ANOVA, two-sided. Statistical significance: *p<0.05, **p<0.01, ***p<0.001, ****p<0.0001. siSLC3A2 is replicated from Fig. 2b. **c**, Volcano plots showing fold-change (log2) versus significance (-log10 p-value) for SLC7A5 versus isotype control. Viral proteins are depicted in orange, SLC7A5 and SLC3A2 – in green. **d**, Maximum intensity projection (MaxIP) of confocal images from A549 cells infected with WSN (Multiplicity of infection (MOI) 3) or mock at 12 hpi. Cells were stained for SLC7A5 (green), M2 (magenta), and nuclei (DAPI, cyan). Scale bars, 20 µm. **e**, Pearson’s correlation coefficient quantification of SLC7A5 and M2 colocalisation in membrane regions. Analysis was performed in mock-infected cells (n = 64) and WSN-infected cells at 12 hpi (n = 119). Statistical significance was determined using the Mann-Whitney test (****p<0.0001), two-sided. **f**, Representative plane and intensity profile showing the co-localisation of SLC7A5 and M2 in A549 cells at 12 hpi. **g,** Cross-linking network of HA. Protein sequences are shown as horizontal bars with structural domains indicated and coloured based on their orientation across the ER membrane. **h,** Early secretory pathway of HA proposed based on the cross-linking and literature data. The % values next to the protein names indicate the impact of the corresponding gene’s knockdown on IAV production using a luciferase assay. The inset boxes show the presumed functions of the respective proteins or maturation steps; the question mark indicates an uncharacterised mechanism. **i,** AlphaFold3 models of glycosylated HA monomers bound to CALR and PDIA3, with cross-links mapped. Each model represents a different glycosylation site, with glycan atoms represented as spheres: N-Acetylglucosamine in blue and Mannose in green. Although the models do not recapitulate HA conformational flexibility and increased oligomeric states during processing, these conceptual models illustrate how cross-links align with CALR-PDIA3 binding to HA glycosylation sites. Cross-links connecting residues within a Cα-Cα distance of ≤30 Å are coloured blue, those between 30–40 Å, a threshold taking protein flexibility into account, are coloured magenta, and those longer than 40 Å are coloured red.

These findings establish SHVIP as a robust approach for identifying host proteins with critical roles in IAV infection. Beyond these examples, infection was affected by the knockdown of several host proteins linked to ER function and paraspeckles, which are discussed in the following sections.

### The steps of the early secretory pathway of HA mapped by cross-linking

We found HA cross-linked to several ER and Golgi apparatus proteins, with all cross-links supporting the same protein orientations across the membrane (Fig. 3g). HA is synthesised in the ER and undergoes folding, N-linked glycosylation, and processing in the ER and Golgi before export to the plasma membrane^55–58^. Examining the ER and Golgi proteins cross-linked to HA indicated that the XL-MS data can be used to reconstruct the sequential steps of the HA secretory pathway along the ER and Golgi compartments (Fig. 3h).

The first step of the pathway is recapitulated by cross-links from HA to calnexin (CANX) and calreticulin (CALR), key chaperones in glycoprotein folding and trafficking^59^, agreeing with the demonstrated role of CANX and CALR in HA processing^60–62^. Cross-links localised to HA stalk and head domains, aligning with AlphaFold3 models of CALR bound to N-glycans on HA (Fig. 3i). HA also cross-linked with protein disulfide isomerase PDIA3, supporting its role in the disulfide bond formation of HA^63,64^, as well as with protein disulfide isomerase TMX1 and an ER chaperone ERP29^65^, not previously linked to HA maturation. Supporting the involvement of PDIA3 and ERP29 in the HA pathway, the knockdown of these proteins in our initial siRNA screen reduced IAV infection (Fig. 2b).

Further along the pathway, HA cross-linked with ERGIC-53 and VIP36 (also known as LMAN1 and LMAN2, respectively, with LMAN2 only detected in the non-stringently filtered cross-link list, Extended Data Table 1e-g), lectins involved in glycoprotein trafficking^66–69^. In support of the role of VIP36 in the HA maturation pathway, VIP36 knockdown significantly reduced IAV infection (Fig. 2b). While ERGIC-53 supports glycoprotein production in other viruses^68^, VIP36 has not yet been implicated in influenza infection. The interaction of ERGIC-53 and VIP36 with HA may facilitate the transport of HA through the Golgi apparatus and possibly aid in its sorting to the plasma membrane. HA also cross-linked with Beta-1,4-galactosyltransferase 1 (B4GALT1), which resides in the Golgi, suggesting B4GALT1 involvement in further processing of HA, and TM9SF2, a protein implicated in Golgi and endosomal sorting^70–74^. This interaction could explain that TM9SF2 overexpression inhibits IAV infection^75^, suggesting a potential antiviral role of TM9SF2.

In summary, these results suggest a pathway in which HA interacts with CANX, CALR, ERP57, ERP29, and TMX1 in the ER, traffics via ERGIC-53 and VIP36, and undergoes further processing by B4GALT1 in the Golgi, possibly regulated by TM9SF2 (Fig. 3h). This analysis demonstrates that in-cell XL-MS can effectively map infection pathways in their native cellular context, capturing sequential protein interactions with high spatial and functional resolution.

### IAV proteins interact with paraspeckles and disrupt their integrity

To determine the subcellular localisation context of the IAV–host protein interactions, we mapped our host proteins crosslinked to viral proteins onto reference spatial proteomics datasets - a global proximity-dependent biotinylation map of human cellular compartments^76^ (Extended Data Fig. 8a). Enrichment analysis revealed a striking overrepresentation of IAV-associated host proteins within nucleolar bodies (Extended Data Fig. 8b). A recently published high-resolution nuclear body proteome^77^ further enabled us to refine this localisation, identifying paraspeckles as a highly enriched subnuclear structure where IAV–host interactions are concentrated (Extended Data Fig. 8c). SHVIP identified cross-links from two viral proteins (NP and NS1) to 14 host proteins that are components of paraspeckles (Fig. 4a). Paraspeckles are membraneless organelles in the nucleus that are built around long non-coding RNA NEAT1_2, which is essential for their structural integrity (Fig. 4b)^78–83^. The DBHS (Drosophila behaviour, human splicing) family proteins – NONO, SFPQ, and PSPC1 – form the paraspeckle core. Additional proteins such as FUS, CPSF6, and HNRNPK contribute to paraspeckle structure and function^78,84–87^. Paraspeckles modulate gene expression and may influence viral replication and pathogenesis through interactions with viral components^78,79,88^.

**Fig. 4.**
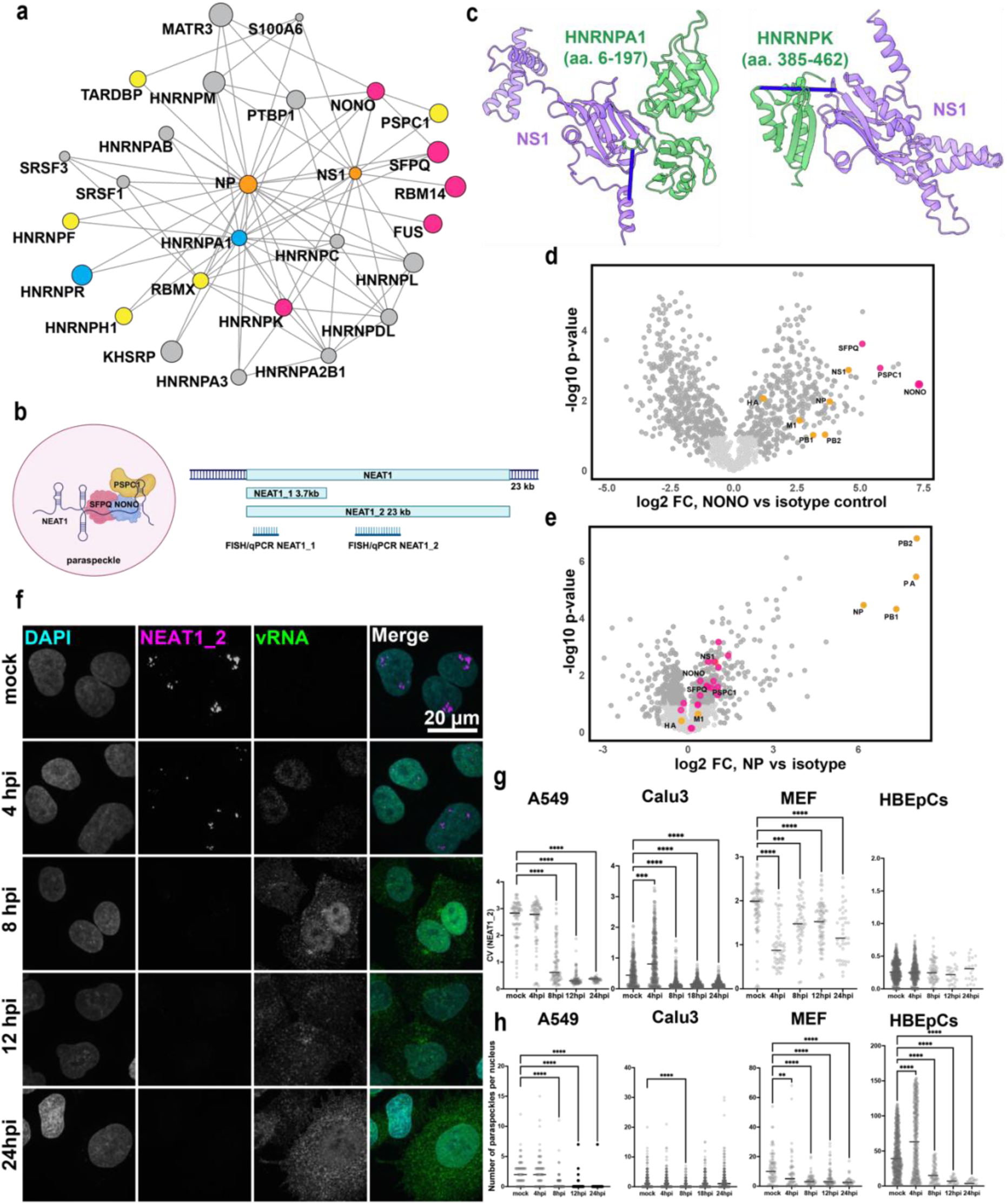
Impact of IAV infection on the paraspeckle formation and NEAT1 expression. **a,** Cross-linking network between paraspeckle and viral proteins. Pink – essential for paraspeckle formation, blue – important, yellow – localised to paraspeckles but dispensable^81^, gray – other proteins cross-linked to both paraspeckle and orange - viral proteins. **b,** Paraspeckle structure, showing the interaction between NEAT1 long noncoding RNA and proteins such as SFPQ and NONO (left) and the NEAT1 isoforms, NEAT1_1 and NEAT1_2, and the position of FISH probes/qPCR primers on NEAT1 used in this study (right). Created in bioRender. https://BioRender.com/b92y974. **c,** AlphaFold models of NS1 and paraspeckle proteins – HNRNPK and HNRNPA1 (See also Extended Data Fig. 6). Cross-links are indicated with blue sticks. **d,** AP-MS of NONO versus IgG controls from WSN-infected A549 cells (14 hpi, n=3). NONO and its known interactors are coloured pink and viral proteins - orange. **e,** AP-MS of NP versus IgG controls from WSN-infected A549 cells (14 hpi, n=3). Proteins that were also identified in SHVIP are highlighted. Proteins coloured as in **d**. **f,** Maximum projection of confocal microscopy images of A549 cells infected with WSN (MOI 3) at 4, 8, and 12 hpi. NEAT1 - magenta, vRNA (PB2 fragment) - green, and DNA (DAPI) - grey. **g,** CV of NEAT1_2 (NEAT1_1 in case of MEF) in the nucleus across different cell lines infected with WSN. **h,** Number of paraspeckles per nucleus across different cell lines infected with WSN. **(g-h),** Each point represents a single nucleus; numbers per condition range from n = 20 to 526. Statistical analysis was performed using ordinary one-way ANOVA with Dunnett’s multiple comparisons test, two-sided. Statistical significance: *p<0.05, **p<0.01, ***p<0.001, ****p<0.0001.

In support of our SHVIP data (Fig. 4a), prior studies have associated the paraspeckle proteins SFPQ, HNRNPK, HNRNPA1, and RBM14 with the IAV infection cycle^5,7,8,89,90^ through interactions with IAV proteins NP and NS1^91–96^. Furthermore, we found that knockdowns of SFPQ and NONO affected IAV infection (Fig. 2b). Well scoring models were obtained for interactions of NS1 with paraspeckle proteins HNRPK and HNRNPA1 (Fig. 4c, Extended Data Fig. 6). We first performed reciprocal immunoprecipitations to confirm interactions of NP with paraspeckle proteins, including NONO, SPFQ and PSPC1 in HEK293T cells (Extended Data Fig. 9a) and WSN-infected A549 cells (Extended Data Fig. 9b-c). Likewise, NONO AP-MS experiment showed enrichment of NONO cross-linking partners such as SPFQ, PSPC1, as well as viral proteins NS1 and NP (Fig. 4d). Similarly, an AP-MS experiment targeting NP showed that proteins cross-linked to NP were overall enriched (Fig. 4e). Taken together, the XL-MS and AP-MS data indicate interactions of IAV NP and NS1 with paraspeckle proteins, particularly the core component NONO. Moreover, the overlap between NP, NS1 and NONO interactors (Extended Data Fig. 9d-e) suggests that other paraspeckle proteins may also play a role in the viral replication cycle during IAV infection.

Next, to assess the effects of IAV infection on paraspeckles, we used fluorescence in-situ hybridisation (FISH) to localise and quantify NEAT1_1 and NEAT1_2 in WSN-infected compared to non-infected A549 cells (Extended Data Fig. 9f and Fig. 4f). While paraspeckles were present in non-infected cells, their number and NEAT1_2 coefficient of variation (CV), which measures signal heterogeneity independently of the mean (see Methods), progressively decreased during infection to very low levels between 4 and 8 hpi (Fig. 4g-h and Extended Data Fig. 9f-g). Moreover, in A549 cells overexpressing mEGFP-NONO and mEGFP-SFPQ, these two proteins co-localised with NEAT1_2 (Extended Data Fig. 9h-k), but redistributed to the nucleoplasm during IAV infection (Extended Data Video 1). Similar reductions in NEAT1_2 CV (Fig. 4g) and paraspeckle numbers (Fig. 4h) were observed in WSN-infected Calu-3 cells (Extended Data Fig. 9l) and mouse embryonic fibroblasts (MEFs) (Extended Data Fig. 9m).

We then extended our analysis to additional IAV strains and primary human bronchial epithelial cells (HBEpCs). The paraspeckle numbers and CV were also reduced in HBEpCs, though with a smaller magnitude, likely due to lower infection efficiency (Extended Data Fig. 9n). H1N1/California/pdm09 (Extended Data Fig. 9o-p) and H3N2/Aichi/68 (Extended Data Fig. 9q-r) strains also reduced NEAT1_2 CV and paraspeckle numbers in A549 and primary HBEpCs cells (Extended Data Fig. 9s-v). Altogether, the results show that IAV disrupts paraspeckles between 4 and 8 hpi consistently across different cell lines, including primary cells, and across different viral strains.

To determine whether and which viral proteins contribute to the paraspeckle disruption, we expressed viral proteins NP and NS1 in HEK293T cells. We showed that the individual expression of either NP or NS1 reduces paraspeckle abundance. Co-expression of vRNPs further intensified paraspeckle dissolution compared to NP alone (Fig. 5a-c), suggesting that, in addition to NP and NS1, other vRNP components may also affect paraspeckle integrity. We hypothesised that the viral endonuclease PA-X may be responsible, as it is produced via frameshift^11^ from the mRNA of the vRNP component PA and its cleavage motif GCUG^11^ is present 139 times in NEAT1_2. Indeed, PA-X overexpression in HEK293T cells significantly reduced paraspeckle number (Extended Data Fig. 9d-f), supporting its role in the maintenance of paraspeckle integrity. Complementarily, we assessed the correlation of NEAT1_2 levels and paraspeckle disassembly by qPCR. NEAT1_2 levels were reduced by 50% at 12 hpi in A549 cells and by 15% at 14 hpi in Calu-3 cells (Fig. 5j). Paraspeckle disruption occurred earlier (4–8 hpi) in both cell lines (Fig. 4g-h, Extended Data Fig. 9l, suggesting that structural disassembly precedes NEAT1 downregulation.

**Fig. 5.**
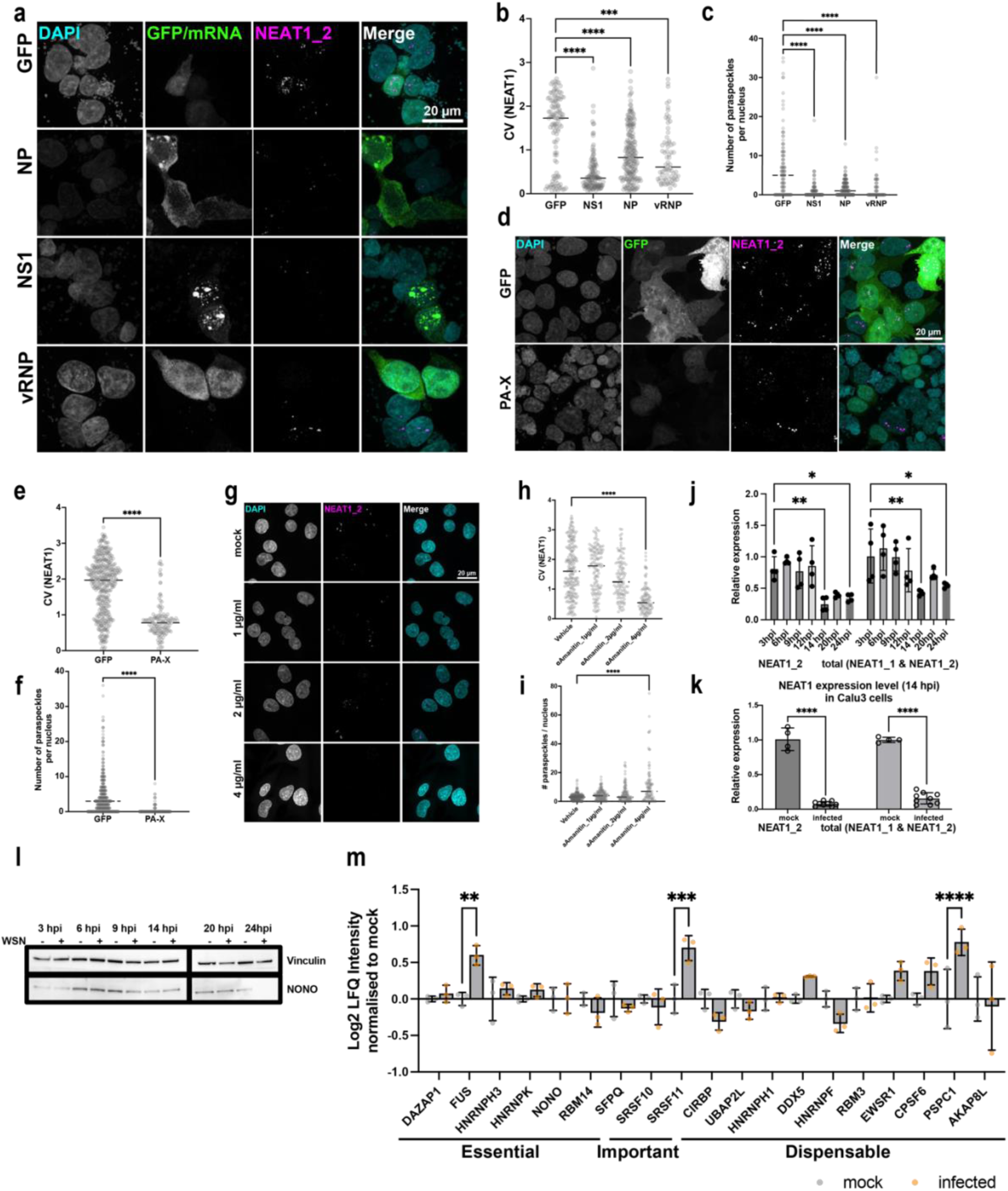
Mechanisms of paraspeckle disruption by IAV proteins. **a,** Maximum projection of FISH confocal microscopy images of HEK293T cells transfected with either GFP (control), WSN viral proteins – NP, NS1, or a mini replicon vRNP system (including PA, PB1, PB2, NP and vRNA of NP), taken at 24 hpi. NEAT1_2 (paraspeckles) is shown in magenta, and GFP (control) or viral mRNAs (NP or NS1) are indicated in green, DNA is counterstained with DAPI (cyan). Scale bar: 20 µm. **b,c,** CV of NEAT1_2 (**b**) and the number of paraspeckles per nucleus (**c**) across different conditions. Only GFP-positive cells were considered. Each point represents a single nucleus; numbers per condition range from n = 72 to 194. Statistical analysis was performed using ordinary one-way ANOVA with Dunnett’s multiple comparisons test, two-sided. Statistical significance: ***p<0.001, ****p<0.0001. **d**, Maximum projection of confocal microscopy images of HEK293T cells transfected with either GFP (control) or co-transfected with PA-X and GFP, taken at 24 hpt. NEAT1_2 (paraspeckles) is shown in magenta, and GFP is indicated in green, DNA is counterstained with DAPI (grey). NEAT1_2 is used as a marker for paraspeckles to assess the impact of IAV proteins on paraspeckle integrity. **e,f,** CV of NEAT1_2 in the nucleus and the number of paraspeckles per nucleus (f) across different conditions. Only GFP-positive cells were considered. Each point represents a single nucleus; numbers per condition range from n = 134 to 399. Statistical analysis was performed using ordinary one-way ANOVA with Dunnett’s multiple comparisons test, two-sided. Statistical significance: ****p<0.0001. **g,** Confocal images showing the maximum projection of NEAT1_2 (magenta) in A549 cells treated with different concentrations of α-Amanitin for 20 h. DAPI (cyan) counterstains the nuclei. Scale bar = 20 μm. h,i CV of NEAT1_2 signal intensity **(h)** and the number of paraspeckles **(i)** per nucleus in A549 cells treated with α-Amanitin for 20 h. Each point represents a single nucleus; numbers per condition range from n = 100 to 207. Statistical analysis was performed using ordinary one-way ANOVA with Dunnett’s multiple comparisons test, two-sided. Statistical significance: ****p<0.0001. **j,** Quantitative PCR analysis of NEAT1_2 and NEAT1_1 expression in A549 cells infected with WSN (MOI 3) normalised to GAPDH and to the mock-infected condition at each corresponding time point (n=3). **k,** Quantitative PCR analysis of NEAT1_2 and NEAT1_1 expression in Calu-3 cells infected with WSN (MOI 3) normalised to GAPDH and to the mock-infected cells (n=3). **l,** Western blot analysis of NONO protein levels in A549 cells infected with WSN (MOI 3) at different time points post-infection (3, 6, 9, 14, 20, and 24 hpi). Vinculin serves as the loading control. **m,** Log₂ Label-Free Quantification (LFQ) intensity of paraspeckle proteins in A549 cells at 14 hpi with WSN (MOI=3), normalised to mock-infected cells (n=3). **(j-k, m)** Data are represented as mean + SD. Statistical analysis was performed using ordinary one-way ANOVA, two-sided. Statistical significance: *p<0.05, **p<0.01, ***p<0.001, ****p<0.0001.

Finally, since paraspeckles depend on an active transcription^84^ and viral polymerase is known to bind and inhibit Pol II^97–99^, we investigated whether Pol II inhibition disrupts paraspeckles. Indeed, treating non-infected A549 cells with Pol II inhibitor alpha-Amanitin^100^ disrupted paraspeckles in a concentration-dependent manner starting 20 hours post-treatment (Fig. 5g-i). Since paraspeckle disruption starts within the first 8 hours of infection, this delay suggests that Pol II inhibition contributes to paraspeckle disruption at later infection stages. NONO levels remained stable between mock and infected A549 cells, indicating that paraspeckle disassembly is not caused by NONO downregulation (Fig. 5l). Consistently, other essential paraspeckle proteins, such as SFPQ and HNRNPK, also showed no significant reduction in expression. Only FUS, SRSF10 and PSPC1 exhibited a significant increase upon infection (Fig. 5m).

Collectively, our findings suggest a three-pronged attack on paraspeckle formation by IAV: NS1- and NP-mediated interactions with paraspeckle protein components, PA-X-dependent degradation of the architectural NEAT1_2 RNA, and possible transcriptional downregulation of NEAT1_2 via Pol II inhibition.

### Disruption of paraspeckles enhances IAV replication by liberating pro-viral host factors

To evaluate the functional relevance of paraspeckle disruption during infection, we used RNA interference to downregulate paraspeckle proteins and measured IAV replication by luciferase assay (Fig. 6A). In agreement with our initial siRNA screen (Fig. 2b), SFPQ knock-down decreased IAV replication in A549 cells, confirming its role in enhancing viral IAV RNA processing^89,90^, whereas NONO knock-down had the opposite effect. In addition, we observed pro-viral effects when knocking down FUS and NEAT1 (Fig. 6a). NEAT1 and NONO knockdown substantially upregulated vRNA, cRNA, and mRNA levels of NP and HA, as shown by strand-specific qPCR^101^ (Fig. 6b, Extended Data Fig. 10a-c). In contrast, SFPQ knockdown downregulated all these viral RNAs, aligning with its role in IAV RNA processing^89,90^. Knockout of NEAT1 and NONO impaired paraspeckle formation and increased IAV replication by 2–2.5-fold (Extended Data Fig. 10f-g, Fig. 6c-d, Extended Data Fig. 10i-k). Overexpressing mEGFP-NONO in wild-type cells impaired IAV replication by 30%, supporting an antiviral role of NONO (Fig. 6e). Supplementing mEGFP-NONO in NONO KO cells reduced viral replication (significantly in one clone and without reaching statistical significance in the second clone, Fig. 6e). Similarly, a minimal NEAT1_2^102^ construct partially rescued paraspeckle formation, whereby a low expression level of NEAT_2 (5 %) likely prevented a more complete rescue (Extended Data Fig. 10h).

**Fig. 6.**
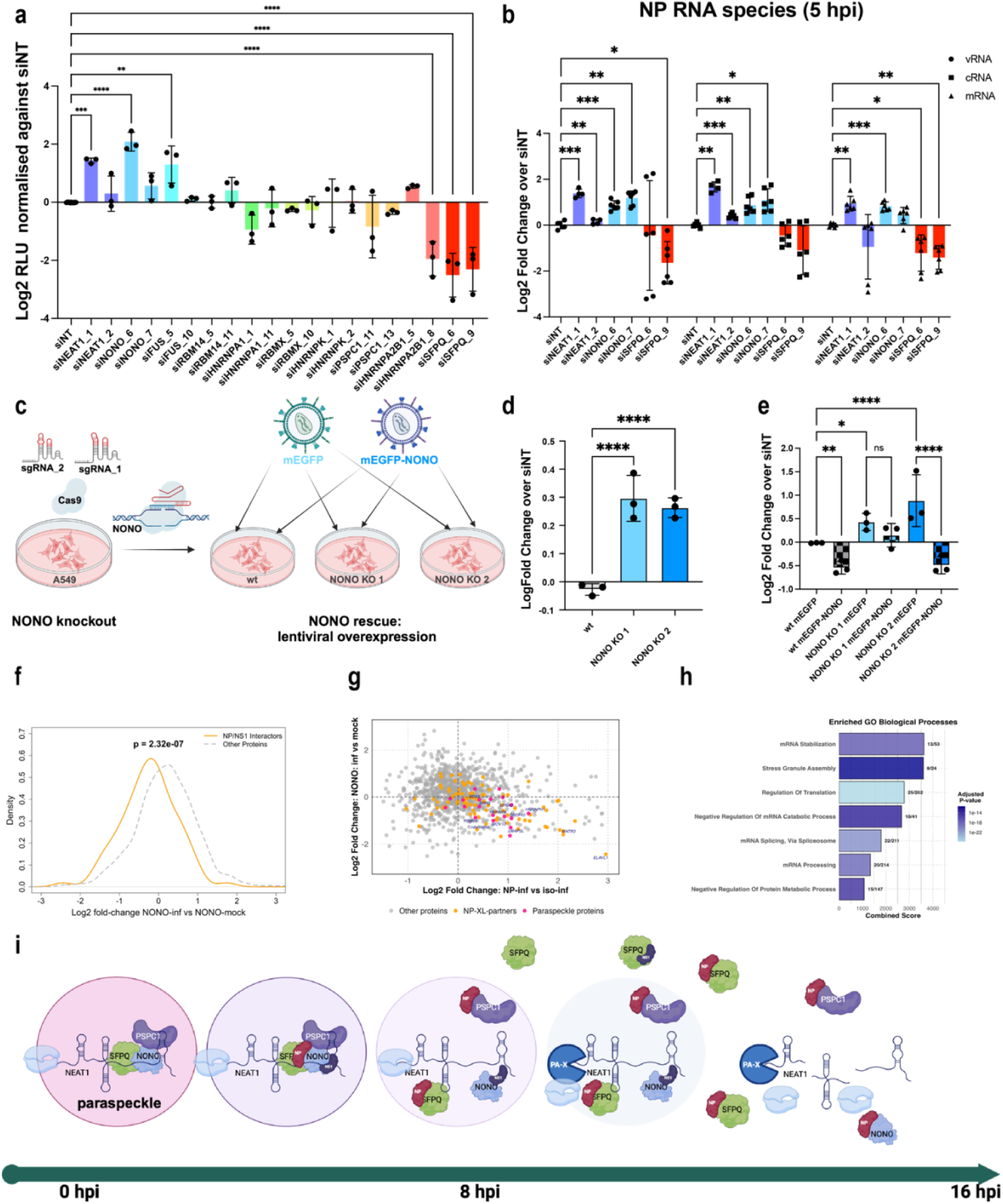
Functional analysis of paraspeckles during IAV infection. **a,** Luciferase activity in A549 cells transfected with siRNAs targeting paraspeckle components. Data are represented as mean ± SD (n = 3). siNEAT1_1 and 2 indicate two different siRNAs both targeting the whole NEAT1.The data for RBM14, RBMX, RBM5, HNRNPA1, HNRNPA2B1, HNRNPK and PSPC1 were reproduced from Extended data Fig. 5c. **b,** Quantitative PCR analysis of viral RNA species of fragment NP normalised to GAPDH in A549 cells with knockdowns of paraspeckle proteins, infected with WSN (MOI 3) at 6hpi. Data are presented as log-fold-change of as mean ± SD relative to the siNT condition (n=3). Mixed effect model with Geisser–Greenhouse correction with Dunnett’s multiple comparisons test was used. **c,** Schematic of NONO knockout and rescue workflow. CRISPR-Cas9 with two sgRNAs generated NONO KO1 and KO2 A549 cell lines. Lentiviral overexpression of mEGFP-NONO or mEGFP (control) in KO cells was used for rescue. Wild-type, NONO KO, and rescued cells were analysed for paraspeckle function and viral replication. Created in BioRender. https://BioRender.com/t41h404. **d,** Luciferase activity in wt, NONO KO1, and KO2 in A549 cells (WSN with PB2-T2A-NanoLuc, MOI 0.01, 48 hpi). Data are represented as mean ± SD (n = 3). **e,** Luciferase activity in wt, NONO KO1, and KO2 A549 cells lentivirally overexpressing mEGFP (control) or mEGFP-NONO (WSN with PB2-T2A-NanoLuc, MOI 0.01, 48 hpi). Data are represented as mean ± SD (n=3). **f,** Density plot of log2-fold-changes of proteins cross-linked to NONO, NP and NS1 identified by AP-MS (NONO infected vs. NONO mock). Proteins cross-linked to NP and NS1 (orange) are co-depleted in infected cells compared to proteins not linked to NP or NS1 (grey). **g,** Log2-fold changes of interactors identified by AP-MS against NONO (infected vs. mock) and NP (infected vs. infected isotype control). Proteins cross-linked to NONO (pink), NP (orange), and non-associated proteins (grey) are shown. **h,** Selected protein categories enriched in both NP and NONO-mock AP-MS datasets from G (see also Table S3). **i,** Schematic representation of paraspeckles disruption: IAV proteins, particularly NP and NS1, disrupt paraspeckle integrity by binding core proteins like SFPQ and NONO, and possibly NEAT1, initiating paraspeckle disassembly. As the infection progresses, PA-X promotes NEAT1 degradation, while POL II inhibition further destabilises paraspeckles, leading to their complete disruption. Created in BioRender. https://BioRender.com/y54j344. **(a-b, d-e)** Data are represented as mean ± SD. Statistical analysis was performed using ordinary one-way (a) or two-way (d-e) ANOVA with Dunnett’s multiple comparisons test, two-sided. Statistical significance: *p<0.05, **p<0.01, ***p<0.001, ****p<0.0001.

To test whether proteins that play roles during IAV infection but are normally sequestered within paraspeckles become available for interaction with IAV proteins upon infection (Extended Data Fig. 10b-c), we compared protein levels in AP-MS targeting NONO in infected cells (disrupted paraspeckles) with non-infected cells (intact paraspeckles). Indeed, proteins cross-linked with NP or NS1 were depleted from the NONO interactome in infected cells (Fig. 6f, Extended Data Fig. 10l-m). Importantly, these changes were not due to infection-induced changes in overall protein levels, as input samples showed no differences between infected and mock-infected cells (Extended Data Fig. 10o-q). This suggests that IAV infection triggers a redistribution of interactors from NONO to viral proteins. In support of this, the key RNA-binding proteins (RBPs) HNRNPC, HNRNPA1, MATR3, SFPQ, and HNRNPK are enriched in AP-MS targeting NP but depleted in AP-MS targeting NONO in infected cells (Fig. 6g-h). Considering that all these RBPs are known regulators of paraspeckle integrity, RNA processing and transcriptional control^86,103–105^, their liberation from NONO and engagement with NP/NS1 may conceivably enhance viral replication. Such a mechanism would be in line with the proviral effects of SFPQ we observed, as well as the documented roles of SFPQ and HNRNPK in IAV RNA splicing and replication^7,8,89,90,106^.

## Discussion

Our in-cell XL-MS using SHVIP provides a high-resolution spatially resolved map of PPIs in IAV-infected cells, revealing host-pathogen contact sites within their native cellular environment. Unlike prior IAV interactome studies based on AP-MS or Y2H, which rely on lysed cells and disrupt native complexes, our approach preserves physiological interactions and captures contacts specific to a late infection stage. For example, the dataset lacks known PPIs from earlier infection stages, e.g. between RNA polymerase II and RdRp^107,108^ as well as between the LRR domains of IAV replication co-factor ANP32 and the RdRp subunit PB2^35,109,110^. Instead, the ANP32A/B LRR domains cross-linked with the C-terminal region of M1. This suggests that, at a late infection stage, M1 may obstruct the ANP32A/B-RdRp interface, potentially inactivating replication and transcription, and thus freeing vRNPs for export from the nucleus. Furthermore, our data enabled us to trace the steps of HA maturation and export through the secretory pathway (Fig. 3c). This exemplifies how in-cell XL-MS can resolve membrane-associated processes with sub-organelle resolution. Beyond implications for understanding the IAV infection cycle, these findings highlight the utility of in-cell XL-MS for reconstructing sequential steps of molecular pathways involved in infection.

Guided by SHVIP, we discovered that IAV actively disrupts paraspeckles, membraneless organelles involved in stress responses, immune regulation, and nuclear protein sequestration^78,81,83^. These phase-separated compartments represent another class of spatially organised cellular structures that can be targeted by viral proteins. We propose that paraspeckle disassembly benefits IAV in two ways. First, it liberates host RNA-binding proteins essential for viral replication. This includes SFPQ, known for its role in IAV RNA expression^90^, as well as HNRNPC and HNRNPA1, which likely facilitate viral RNA synthesis and nuclear transport^92,111,112^. Second, paraspeckle disassembly may suppress antiviral responses. Given their roles in stress adaptation and immune signalling, and our findings that NEAT1 and NONO restrict viral replication/transcription, the loss of paraspeckles could attenuate host antiviral gene regulation^113,114^. Thus, we posit that paraspeckle dismantling is an active viral strategy rather than a secondary consequence of host transcriptional repression.

Interestingly, while paraspeckles disappeared in IAV-infected A549 and Calu-3 cells, they persisted in HeLa cells infected with IAV, according to a previous study^89^. HeLa cells are less permissive to IAV^115–119^, and this may reflect an inability of the virus to counteract paraspeckle-mediated antiviral functions in restricted infections. Moreover, our data on NEAT1_2 depletion align with evidence showing increased non-canonical splicing of the long isoform (NEAT1_2) during IAV infection, possibly due to NEAT1_2 release from paraspeckles, increasing its availability for splicing^120^. Notably, other viruses, such as Hantavirus^121^, HSV-1^89,122^, and Dengue virus^123^, induce NEAT1 expression and enhance paraspeckle formation rather than disrupting them. However, for IAV, the diffused paraspeckle staining and weaker NEAT1_2 upregulation compared to HSV-1 were noted^120^. This may reflect a fundamental difference in viral countermeasures: IAV actively dismantles paraspeckles to hijack their protein components, while other viruses may tolerate or exploit them for antiviral signalling.

While validating all PPIs and contact sites in our map is beyond the scope of a single study, our snapshot provides a foundation for numerous follow-up investigations. For instance, we found five viral proteins, including all three polymerase subunits, cross-linked with calcium homeostasis-related proteins. Individual knockdowns did not significantly affect infection (Extended data Fig. 5), likely due to functional redundancy. However, these interactions align with previous reports linking calcium levels to influenza virus infection^4,60,124^, supporting a novel hypothesis that calcium-mediated effects on infection involve viral interactions with calcium-binding proteins.

In the future, in-cell XL-MS of IAV-host interactions could be extended to other IAV strains and to earlier infection stages to capture the dynamics of these interactions, including those involving IAV RdRp during transcription^107,109^. Although AlphaFold has limitations in modelling host-pathogen interactions, and only a subset of PPIs could be modelled here, future advancements in structural prediction—particularly those integrating cross-links as restraints—could use XL-MS data to provide a detailed map of the structural interface between IAV and human proteins.

In conclusion, our work provides a vast and unique resource of native IAV-host PPIs and serves as a blueprint demonstrating how integrating in-cell XL-MS, functional analyses, and structural modelling can resolve the native spatial and structural organisation of viral-host interfaces and uncover mechanisms of viral subversion and host cell perturbation.

## Methods

### Immortalised cell lines

Human A549 (ECACC 86012804) and HEK293T (ECACC 120220101) cells were cultured in DMEM (Gibco 11960044) supplemented with 10% FBS (Gibco 0270106), GlutaMAX (Gibco 35050038), sodium pyruvate (Gibco 11360039), and penicillin– streptomycin (Gibco 15070063) at 37 °C and 5% CO₂. Calu-3 cells (gift from Petr Chlanda) were maintained in the same medium with 20% FBS. MDCK (ECACC 84121903), MDCK.II (ECACC 00062107), and MDBK cells (gift from Ervin Fodor) were grown in MEM supplemented with 10% FBS, GlutaMAX, sodium pyruvate, and penicillin–streptomycin. MEFs (gift from G. Gabriel) were cultured in DMEM with 10% FBS, GlutaMAX, sodium pyruvate, penicillin–streptomycin, and non-essential amino acids (Gibco 11140050).

### Primary cells

Primary cells HBEpC (Promocell C-12640, donor lot #499Z012.1) were maintained in Airway Epithelial Cell Growth Medium (Promocell C-21060) were cultured according to the manufacturer’s instructions in Airway Epithelial Cell Growth Medium (Promocell, C-21060) at 37 °C with 5% CO₂. Primary cells were used at passages 2–6 for all experiments. All cell lines were routinely tested for mycoplasma contamination (MycoplasmaCheck, Eurofins).

### Virus production

WSN was produced in confluent MDBK cells infected at a multiplicity of infection (MOI) of 0.001 in DMEM containing 0.5% FBS. After 36 h incubation at 37 °C and 5% CO₂, when full cytopathic effect (CPE) was observed, supernatants were harvested, clarified by centrifugation (1,500 g, 10 min, 4 °C), and aliquoted for storage at −80 °C. pdm09 and H3N2 were produced in MDCK cells infected at an MOI of 0.001 in serum-free DMEM supplemented with 0.2% BSA and 1 µg/mL TPCK-treated trypsin. After 36–48 h incubation at 37 °C and 5% CO₂ and confirmation of full CPE and successful HA assay, viral supernatants were clarified and stored as above.

### Generation of Recombinant IAV Overexpressing NanoLuc on PB2

A recombinant influenza A virus expressing NanoLuc fused to PB2 was generated using a 12-plasmid reverse genetics system as described before^50^. The system included eight plasmids encoding the viral RNA segments and four expression plasmids for PB2, PB1, PA, and NP under control of a human Pol I promoter. The NanoLuc reporter gene was inserted in-frame at the N-terminus of PB2, separated by a self-cleaving T2A peptide to allow independent translation.

### HA assay

The HA assay was performed in 96-well round-bottom plates using two-fold serial dilutions of virus in PBS (50 µL per well). An equal volume (50 µL) of 0.5% chicken erythrocyte suspension was added to each well. Plates were incubated for 30 min at 4 °C, followed by 30 min at room temperature. HA titres were determined visually as the highest virus dilution producing complete haemagglutination, defined by a uniform lattice across the well. Results are expressed in haemagglutination units (HAU).

### Plaque Assay

For WSN, confluent MDCK cells in 6-well plates were infected with 330 µL of serially diluted virus in infection media (DMEM with 0.2% BSA). Plates were incubated for 30 min at 4 °C, followed by 30 min at 37 °C with gentle rocking. Cells were overlaid with avicel-containing media (DMEM, 0.2% BSA, avicel) and incubated for 72 h. After removing the overlay, cells were washed with PBS and stained with 1% crystal violet for 5 min. Plaques were counted to calculate viral titre in PFU/mL. For pdm09 and H3N2, MDCK-II cells were infected as above and overlaid with DMEM containing avicel, 0.2% BSA, and 1 µg/mL TPCK-treated trypsin. After 72 h, cells were fixed with 4% paraformaldehyde for 30 min at 4 °C and blocked with 0.3% hydrogen peroxide. Viral plaques were detected by immunostaining with anti-NP antibodies (Aichi: ThermoFisher MA5-42364; pdm09: Abcam ab43821; 1:1000) for 1 h at room temperature, followed by HRP-conjugated secondary antibody and KPL TrueBlue substrate (15 min). Viral titres were calculated as PFU/mL:

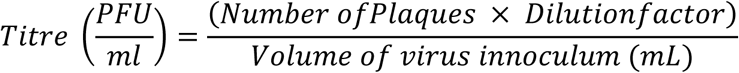

### Luciferase assay

A549 cells were seeded at 1 × 10^5^ cells per well in 24-well plates and transfected 12 h later in duplicate with 10 µM siRNAs using RNAiMAX (Thermo Fisher Scientific), according to the manufacturer’s instructions. After 72 h incubation at 37 °C and 5% CO₂, cells were washed with infection media and infected with PB2-T2A-NanoLuc recombinant WSN at an MOI of 0.01. At 48 h post-infection, cells were lysed in 200 µL Passive Lysis Buffer (Promega) per well and incubated at room temperature for 20– 30 min on a rocking platform. For luciferase activity, 25 µL of lysate was mixed with 25 µL NanoGlo substrate (Promega) in a white 96-well plate. Luminescence was measured after 5–10 min equilibration using a luminometer (1000 ms integration time).

### FISH

Cells (1 × 10^5^ per well) were seeded onto sterilised glass coverslips in 24-well plates and grown to ~70–80% confluence. Cells were washed with DPBS, fixed in 4% paraformaldehyde for 10 min at room temperature, and permeabilised in 70% ethanol overnight at −20 °C. Hybridisation and amplification were performed using the Hybridization Chain Reaction (HCR v3.0) protocol adapted from^125^, with optimisations for detecting IAV vRNA, mRNA, and cRNA (see Supplementary Methods for probe sequences and design). Briefly, fixed cells were pre-hybridised with Probe Hybridization Buffer (Molecular Instruments) and incubated overnight with 2 pmol of each probe at 37 °C. Following washes in Probe Wash Buffer and SSCT, amplification was carried out overnight at room temperature using snap-cooled HCR hairpins (h1 and h2). Cells were washed in 5× SSCT, stained with DAPI, and mounted using Dako antifade mountant (Agilent). Samples were imaged using a confocal microscope with appropriate filter settings for DAPI and HCR fluorophores. Acquisition settings were optimised to reduce background and enhance signal detection.

### siRNA transfection

All siRNAs were purchased from Qiagen (see Supplementary Table). A549 cells at ~60% confluence were transfected with 5 pmol siRNA using Lipofectamine RNAiMAX (Thermo Fisher Scientific, 13778075) according to the manufacturer’s protocol. Knockdown efficiency was assessed 72 h post-transfection by qPCR or western blot.

### Cell viability

Cell viability was measured using the CellTiter-Glo® 2.0 Assay Kit (Promega, G9241). Treated cells in 24-well plates were incubated with reagent directly added to the culture medium, followed by 10 min shaking at room temperature. Luminescence, proportional to intracellular ATP levels, was measured using a plate reader and normalised to untreated controls.

### CRISPR/Cas 9-mediated knock-out cell lines

sgRNAs were designed using Benchling or purchased as validated guides (IDT) and resuspended at 1 µM. Recombinant Cas9 protein (EMBL Protein Production Facility) was diluted to 1 µM in Opti-MEM. A549 cells were seeded at 1.5 × 10^5^ cells/well in 6-well plates at <50% confluence to ensure high knockout efficiency. Cas9–sgRNA ribonucleoprotein (RNP) complexes were formed in Opti-MEM and transfected using Lipofectamine RNAiMAX. After 96 h, cells were either harvested for genomic DNA (NEB kit) or seeded at 0.5 cells/well in 96-well plates for clonal selection. Target loci were PCR-amplified, gel-purified, and sequenced (Sanger). Indel frequency was analysed using TIDE^121^, and knockout was confirmed by western blotting.

### Plasmid transfection

Plasmid transfections were performed in HEK293T cells at 70–80% confluence using Lipofectamine 3000 (Thermo Fisher Scientific) according to the manufacturer’s instructions. Plasmid DNA and Lipofectamine 3000 reagent were each diluted in Opti-MEM (Thermo Fisher Scientific), combined, and incubated for 15 min at room temperature to allow DNA–lipid complex formation. The complexes were added dropwise to the cells and incubated for 4–6 h at 37 °C in 5% CO₂, after which the medium was replaced with fresh complete DMEM. Cells were incubated for an additional 16–48 h prior to analysis. Transfection efficiency was assessed by co-transfection with a GFP-expressing plasmid or by fluorescence microscopy following immunofluorescence (IF) or FISH staining.

### Lentiviral cell line generation

Lentiviral vectors encoding mEGFP-tagged NONO, PSPC1, or SFPQ were co-transfected into HEK293T cells with packaging plasmids using a calcium phosphate transfection protocol. Supernatants containing lentiviral particles were collected after 48 h, filtered, and stored at −70 °C. A549 cells were transduced with viral supernatants in the presence of polybrene and incubated for 48–72 h to allow stable genomic integration. Transduced cells were selected using 10 µg/mL Blasticidin and maintained in complete DMEM. Expression of tagged proteins was confirmed by fluorescence microscopy and western blotting. Protein functionality was verified by their co-localisation with NEAT1 in paraspeckles.

### RNA Extraction, Reverse Transcription and qPCR

Total RNA was extracted using the RNeasy Kit (Qiagen) according to the manufacturer’s instructions. Cells were lysed in buffer RLT supplemented with 5 mM TCEP, and RNA was purified on RNeasy columns with RW1 and RPE wash buffers. RNA was eluted in RNase-free water, quantified using a NanoDrop spectrophotometer, and stored at −70 °C.

Reverse transcription was performed with up to 2 µg RNA per 10 µL reaction using the High-Capacity cDNA Reverse Transcription Kit (Thermo Fisher Scientific, 4368814). qPCR was carried out using PowerUp SYBR Green Master Mix (Thermo Fisher Scientific, A25742) with gene-specific primers (see Supplementary Table). Thermal cycling and melt curve analysis were performed according to the manufacturer’s protocol. Relative gene expression was quantified using the ΔΔCt method and normalised to GAPDH.

### Immunoblotting

Whole-cell lysates were resolved by SDS–PAGE using 4–12% precast TG PRiME™ gels (SERVA) and transferred to nitrocellulose membranes (Bio-Rad). Membranes were blocked in 2.5% BSA (in TBS with 0.01% Tween-20) for 1 h at room temperature, followed by overnight incubation at 4 °C with primary antibodies diluted in blocking buffer. After three washes in TBS-Tween, membranes were incubated for 1 h at room temperature with HRP-conjugated anti-mouse or anti-rabbit secondary antibodies (1:5,000), washed again, and developed using enhanced chemiluminescence (Thermo Fisher Scientific, A38555). Antibodies used for immunoblotting are listed in Supplementary Table. Uncropped blots are available in the Source Data files.

### Affinity Pulldown of TwinStrep-tagged Proteins

HEK293T cells were transfected with TwinStrep-tagged constructs (NONO, M2, or empty vector control) using PEI at a 1:3 DNA:PEI ratio. At 24 h post-transfection, cells were harvested by scraping in PBS, washed, and lysed in IP lysis buffer (Thermo Fisher) supplemented with HALT protease inhibitor cocktail. Lysates were clarified (17,000 g, 20 min, 4 °C) and incubated with Strep-Tactin magnetic beads (IBA Lifesciences) pre-equilibrated in Buffer W. After 1 h incubation at room temperature with rotation, beads were washed, and bound proteins were eluted in Laemmli buffer at 95 °C for 2 min. Eluates were analysed by SDS–PAGE and Western blot.

### Immunoprecipitation (IP)

A549 cells were seeded in 10 cm dishes (4 × 10⁶ cells/dish) and infected with WSN (MOI 3) for 14 h. Cells were harvested and lysed in IP buffer (Thermo Fisher) supplemented with HALT protease inhibitor (ThermoFisher). For certain IPs (e.g., NP, NONO, SFPQ, PSPC1), 0.1% SDS was added to reduce non-specific binding. Lysates were cleared by centrifugation (max speed, 20 min, 4 °C). IP was performed with 10 µg of antibody and 500 µL of cleared lysate for 4 h at 4 °C, followed by incubation with Protein A/G or G Dynabeads (Thermo Fisher) for 1 h at room temperature. Isotype-matched control antibodies were used in all experiments. Beads were washed with TBS + 0.05% Tween-20 and lysis buffer and finally with TBS alone. For AP-MS, bound proteins were eluted in 8 M guanidine HCl at 95 °C and precipitated in cold ethanol overnight. Inputs were processed by methanol/chloroform precipitation. For Western blotting, elution was performed with 0.2 M glycine (pH 2.0), followed by Tris neutralisation and/or SDS loading buffer. Eluates were analysed by SDS–PAGE, Western blot, or mass spectrometry as described below. All AP-MS experiments were performed in triplicate.

### SHVIP sample preparation

Cells were infected at an MOI of 5 and, at 5 hpi, incubated in labelling medium containing 500 µM L-HPG in methionine-free DMEM with 0.2% BSA. At 14 hpi, cells were washed with PBS and harvested by scraping. For cross-linking, cell pellets were resuspended in PBS and cross-linked with 5 mM DSSO for 1 h at room temperature with constant agitation. The reaction was quenched with 30 mM Tris (pH 8.0) for 20 min. Cells were lysed in buffer containing 200 mM Tris (pH 8.0), 4% CHAPS, 1 M NaCl, 8 M urea, and protease inhibitors. Lysates were frozen at −80 °C until enrichment. After thawing, lysates were treated with Benzonase and sonicated (Bioruptor Pico, 10 × 30 s on/off cycles), then clarified by centrifugation (10,000 g, 10 min). A 50 µL aliquot was saved as input. Enrichment of of HPG-labelled proteins was performed as described before ^29^: 800 µL of lysate was incubated with 200 µL picolyl-azide agarose beads (Click Chemistry Tools), and copper(I)-catalysed click chemistry was performed overnight at room temperature with rotation. Proteins were reduced with 10 mM DTT (70 °C, 15 min) and alkylated with 40 mM chloroacetamide (room temperature, dark). Beads were washed sequentially in gravity columns with: 1% SDS with 250 mM NaCl, 5 mM EDTA in 100 mM Tris (pH 8.0), 8 M urea in 100 mM Tris (pH 8.0), 80% acetonitrile (ACN) in water, 5% ACN in 50 mM TEAB, 5% ACN with 2 M urea in 50 mM TEAB. Bound proteins were digested on-bead overnight at 37 °C with trypsin and LysC in the final wash buffer. Peptides were collected, acidified with 1% formic acid, desalted using C18 stage tips^126^ (or Sep-Pak C8 columns for cross-linked peptides), dried by vacuum centrifugation, and stored at −20 °C until LC–MS or further processing. Input samples were processed by methanol–chloroform precipitation^127^ and digested in buffer containing 1% SDC, 5 mM TCEP, and 40 mM CAA in 50 mM TEAB. Resulting peptides were desalted as above.

### Off-line fractionation of cross-linked peptide samples

To enrich cross-linked peptides, HPG-enriched samples were first subjected to strong cation exchange (SCX) chromatography using a PolySULFETHYL A column. Peptides were separated with a 95-minute linear gradient and collected in 45-second intervals. Fractions were desalted, dried, and stored at −20 °C. After initial measurement, selected SCX fractions were pooled and further separated by size exclusion chromatography (SEC) using a Superdex 30 column. Fractions were collected every 2 min, dried, and stored at −20 °C prior to LC–MS/MS.

### LC-MS/MS of bottom-up proteomics and AP-MS samples

Bottom-up samples and AP-MS eluates were analysed using Orbitrap Fusion Tribrid or Exploris 480 instruments coupled to nano-LC systems. Peptides were loaded onto a 50 cm in-house packed C18 column and separated by 120–180 min gradients.

For bottom-up and AP-MS samples: MS1 scans were acquired at 120,000 resolution, dynamic exclusion was set to 40 s, precursors (charge +2 to +4) were isolated (1.6 m/z window) and fragmented by HCD at 30% normalised collision energy (NCE), MS2 detection was performed in the ion trap (Fusion) or Orbitrap (Exploris).

### LC-MS/MS of cross-linked samples

Cross-linked peptides were analysed on an Orbitrap Fusion Lumos equipped with a FAIMS Pro Duo interface. MS1 scans were performed at 120,000 resolution using FAIMS voltages (−50, −60, −75 V). Precursors (charge +4 to +8) were fragmented by stepped-HCD (21%, 27%, 33% NCE) and MS2 scans acquired in the Orbitrap at 60,000 resolution.

### Bottom-up proteomics data analysis

Raw files were searched using MaxQuant v1.6.2.6a^128^ against the human SwissProt database (release 2020) and the IAV/WSN/1933 (H1N1) proteome. Trypsin was used for in silico digestion (max 2 missed cleavages). Carbamidomethylation (C) was set as fixed modification and variable modifications included, oxidation (M), acetylation (N-term), and methionine replacement with HPG, where appropriate. iBAQ and LFQ quantification were enabled. Contaminants, reverse hits, and proteins identified only by site were removed. For LFQ, missing values were imputed as described before^129^. Fold changes were log2-transformed and p-values were computed using two-sided t-tests. All AP-MS experiments were performed in triplicates.

Raw data were searched using Scout (v1.5.1)^130^ with DSSO cross-linker settings and residue specificity for K, S, T, and Y. Search parameters included: Minimum peptide length: 6, Max missed cleavages: 3, Precursor mass tolerance: 10 ppm, Fragment mass tolerance: 20 ppm, Static mod: carbamidomethylation (C); variable mod: oxidation (M) Data were filtered at 2% or 5% false discovery rate (FDR) on crosslink spectrum match (CSM), residue pair, and protein-protein interaction (PPI) levels. For performance evaluation (Extended Data Fig. 1M-R), SCX fractions were searched per replicate. For the full dataset, SCX and SEC files were combined for global analysis.

### Confocal microscopy

All imaging was conducted using a Nikon Ti2 microscope equipped with a CSU-W1 spinning disk confocal unit and an Andor DU-888 X-11633 camera. The microscope was controlled by NIS-Elements software and utilized a 100x/1.49 SR Apo TIRF AC oil immersion objective (Nikon). For live-cell imaging, cells were seeded in Ibidi μ-Slide 4-well chambers (Ibidi USA, NC0685967) and maintained at 37°C with 5% CO₂. Imaging was performed using the 405 nm (DAPI channel), 488 nm (GFP channel), and 561 nm (Cy5 channel) lasers, with exposure times optimised for each experiment. The camera was maintained at −69.4°C, with a binning of 1×1 and a readout speed of 30 MHz. Z-stacks were captured with an interval size of 130.6 nm, and images were acquired at a resolution of 1024×1024 pixels.

### Image Analysis

All images were processed as maximum intensity projections (MaxIP). Nuclei (DAPI) and paraspeckles (NEAT1_2) were segmented using CellProfiler with the Otsu thresholding method. The coefficient of variation (CV) of the NEAT1_2 signal within nuclei and the number of segmented paraspeckles were used as independent metrics. Both metrics have their respective advantages and disadvantages. While CV relies on more reliable nuclei segmentation, it is an indirect measure of paraspeckle integrity and does not provide precise structural information. In contrast, paraspeckle quantification directly assesses paraspeckle formation but is more error-prone, particularly at later time points when the NEAT1_2 signal diminishes.

### Subcellular localisation analysis of viral protein interactors

To compare subcellular localisations of host interactors across studies, we compiled viral–host protein interactions from SHVIP (this study), Watanabe & Kawakami et al. (2014)^19^, Haas et al. (2023)^21^, and the meta-analysis by Chua et al. (2022)^40^. For all datasets, only viral proteins that were identified in SHVIP (PB1, PB2, NP, NS1, M1, M2, HA, NA) were considered.

Host protein localisations were annotated based on data from the Human Protein Atlas (HPA). Only primary localisation assignments were retained. A simplified set of localisation categories was defined that grouped terms into: Plasma membrane, Cytosol, Cytoskeleton, Vesicular system, Endoplasmic reticulum (ER), Mitochondrion, Nucleoplasm, Nucleoli, Nucleus (other), and Other, followed by manual curation. For ambiguous or multi-localised proteins, the most specific localisation according to UniProt was assigned.

For SHVIP, we included only high-confidence host interactors filtered at 2% FDR at both the crosslinking PPI levels. For Watanabe et al., host interactors were taken directly from their published AP-MS dataset in HEK293 cells expressing individual FLAG-tagged viral proteins. For Haas et al., we used the high-confidence filtered dataset provided by the authors, based on AP-MS of 13 IAV proteins expressed in three cell types across three different IAV strains. For Chua et al., only host interactors identified in at least three independent studies or databases were considered, as described in their meta-analysis^40^.

Viral protein localisation was manually curated from published literature^2,16,38,41,42,51,55,56,137–145^ and database annotations (UniProt). Where conflicting evidence for localisation existed, fractional assignments were split evenly across compartments (e.g., a score of 1 distributed as 0.5 and 0.5 across two compartments).

### Validation of cross-links by host structures and AlphaFold 2 models

For each cross-linked human protein, all structures for this protein were retrieved from the Protein Data Bank (PDB) based on the corresponding UniProt ID using both UniProt^131^ website REST API and RCSB PDB Search API^132^. All cross-linked pairs were mapped to the retrieved structures, and if the corresponding pair was found among structures, Cα-Cα was calculated. If one pair was mapped to multiple structures, the shortest Cα-Cα distance was taken. The same mapping procedure was used for the random cross-linked pairs generated by taking arbitrary lysine-lysine pairs for individual proteins and protein pairs detected in PPIs. AlphaFold 2^44^ models were built using AlphaPulldown^133^. Structural figures were rendered using UCSF ChimeraX^134^.

### Virus-host interaction structural modelling

AlphaFold 2 models were built using AlphaPulldown^133^ version 2.0 using 24 recycles. The sequence and template databases as of August 2024 were used. AlphaFold 3 was run using the original AlphaFold 3 code and using the default sequence database (downloaded in November 2024) and PDB database from November 2021 using the download script provided in AlphaFold 3^48^. AF3x^47,49^ was run with default settings and using version #dfb94a3 and the same databases as AlphaFold 3^48^. The quality of models was assessed using interface predicted template modelling (ipTM) score, predicted template modelling (pTM), residue’s predicted local distance difference test (pLDDT) score, and predicted alignment error (PAE), as returned by all the above tools. Structural figures were rendered using UCSF ChimeraX^134^.

### Other bioinformatics analyses

Functional enrichment analysis was performed using the Enrichr web platform^135^. The default Fisher’s exact test with the Benjamini-Hochberg method for correction for multiple hypotheses testing was used. For cross-links to viral proteins where alternative proteins with identical cross-linked peptide sequences were detected ambiguously (e.g., histone proteins), only the first protein from the list of ambiguous candidates was selected for the analysis. Cross-link diagrams were drawn using xiNET^136^.

## Data availability

Mass spectrometry data will be deposited in the PRIDE database. Structural models will be made available on Zenodo upon publication. Fluorescent images will be deposited in the BioImage Archive.

## Code availability

Not applicable.

## Acknowledgements

DY and JK have been supported by the German Research Foundation (DFG) grant ID: KO 5979/2. KG and JK were supported by DFG VISION grant ID: GRK 2887/1. JK was supported by ERC (TransFORM, 101119142). F.L and L.M have been supported by DFG Project LI 3260/6-1 and F.L was supported the Leibniz-Wettbewerb (P70/2018). B.B. acknowledges funding from Deutsche Forschungsgemeinschaft (https://www.dfg.de/en/index.jsp) grant BO 5917/1-1. This project received funding from the FEBS Excellence Award, of the Deutsche Forschungsgemeinschaft (DFG project number 506356718/ DU 2279/2-1) and the European Molecular Biology Laboratory to O.D. We thank Dr. Ying Wang for her support in the lab, Valentin Mauer for assistance with statistical analysis, and Dr. Lucas DeFelipe for guidance on figure preparation and adherence to publication standards. We thank Caroline Demeret and Nadia Naffakh for critical reading and helpful comments on the manuscript.

## Author contributions

I.K., F.L., B.B., and J.K. conceived and designed the study. I.K., L.M., and B.B. performed experiments, K.G. performed structural analyses and modelling, and created diagram figures, D.Y. contributed to structural modelling, D.Z. helped with data analysis, K.B performed experiments, O.D. supervised experiments, S.S.B and S.B performed experiments, G.B. supervised experiments. I.K., L.M., B.B., K.G, and J.K. analysed data. I.K., B.B., K.G., and J.K. wrote the manuscript. All authors revised the manuscript.

## Declaration of generative AI and AI-assisted technologies

During the preparation of this work, the authors used ChatGPT and Grammarly to improve the style and grammar of the text. After using this tool or service, the authors reviewed and edited the content as needed and take full responsibility for the content of the publication.

## Supplemental Information

**Extended Data Fig. 1.**
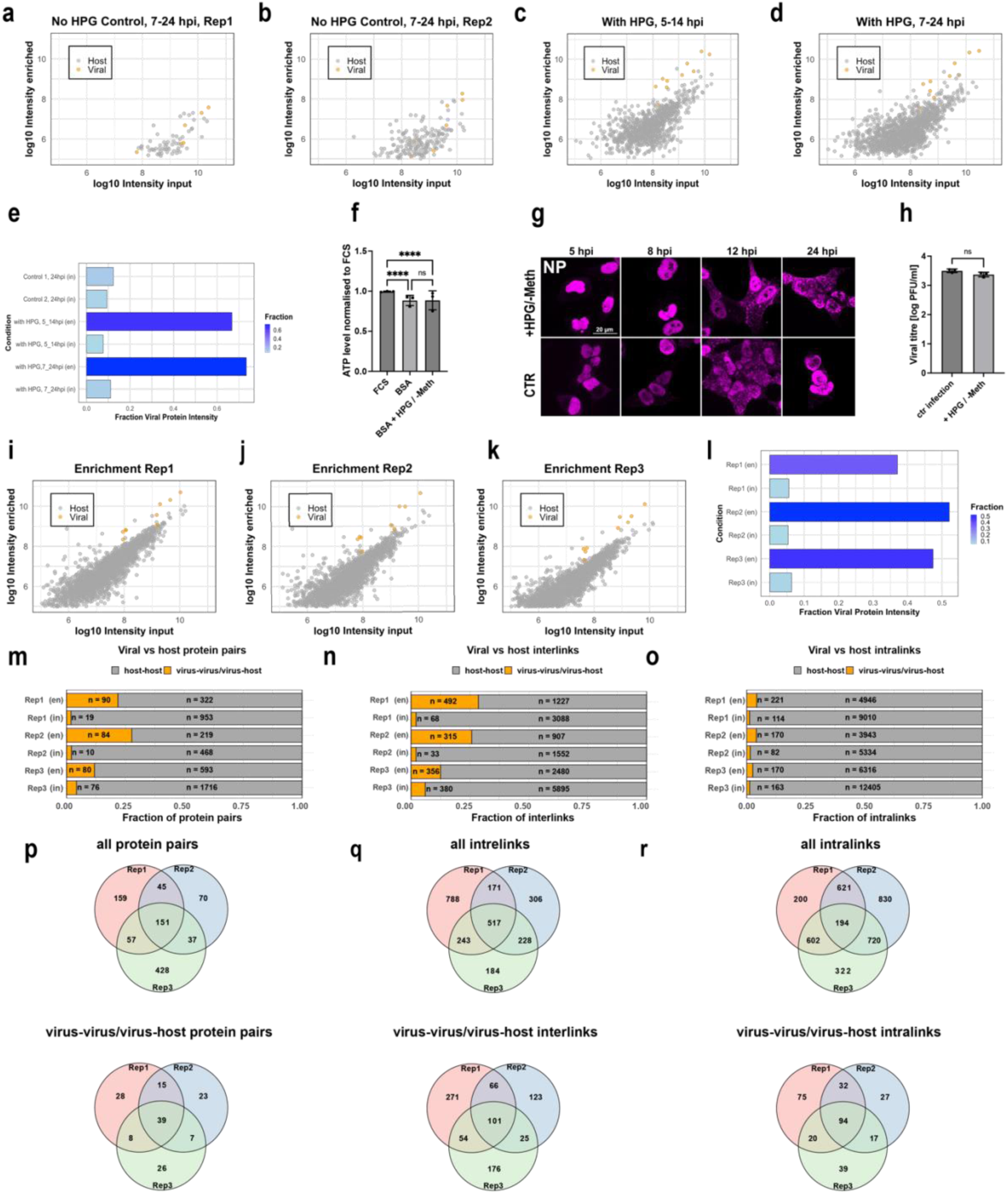
Optimisation of SHVIP and enrichment of IAV proteome in the SHVIP experiment. a-d, Determination of optimal time frames for selective enrichment of viral proteins. Scatter plots represent the correlation between input (x-axis) and enriched (y-axis) log10 iBAQ-values (intensity-based absolute quantification of proteins) of host (grey) and viral (orange) proteins at various time points and conditions. Conditions include no-HPG controls (**a-b**) at 7–24 hpi and HPG-containing samples at 5–14 hpi (**c**) and 7–24 hpi (**d**). L-HPG stands for L-homopropargylglycine. **e,** Fraction of viral protein intensities in total proteomic data under various conditions summarising the data show in **(a-d)**. Bars represent the proportion of viral proteins enriched relative to input proteins across different experimental setups. Enriched samples indicated as en, and input as in. **f,** Viability of A549 cells after incubation in three different media conditions: complete media (FCS), infectious media (BSA), and SHVIP media (BSA L-HPG). Cell viability is normalised to the complete media condition (FCS) (n=3). Data represent mean ± SD. Statistical significance was determined using an ordinary one-way ANOVA. Significance levels are indicated as follows: **** for p < 0.0001 and ns for not significant. **g,** Maximum projection of A549 cells infected with WSN at various time points. The viral nucleoprotein (NP) is used as an infection marker (magenta). Cells were treated with either +HPG/-Meth media or standard infectious media (CTR). Meth stands for methionine. **h,** Viral titres at 14 hpi comparing standard infectious media and +HPG/-Meth media protocols. Data represent mean ± SD (n=3). Statistical significance was determined using an ordinary one-way ANOVA. Significance levels are indicated as follows: ns for not significant. **i-k,** Scatter plots represent the correlation between input (x-axis) and enriched (y-axis) log10 iBAQ-values of host (grey) and viral (orange) proteins for three independent replicates. **l,** Fraction of viral protein intensities in total proteomic data in three replicates summarising the data shown in I-K. Bars represent the proportion of viral proteins enriched relative to input proteins across different experimental setups. Enriched samples are indicated as en, and input as in. **m,** Fraction of viral-viral/viral-host and host-host protein pairs across three replicates. **n,** Fraction of viral-viral/viral-host and host-host interlinks across three replicates. **o**, Fraction of viral-viral/viral-host and host-host intralinks across three replicates. **p,** Venn-diagram depicting overlap of all protein pairs (above) and viral-viral/viral host protein pairs among three replicates. **q,** Venn-diagram depicting overlap of all interlinks (above) and viral-viral/viral host interlinks among three replicates. **r,** Venn-diagram depicting overlap of all intralinks (above) and viral-viral/viral host intralinks among three replicates.

**Extended Data Fig. 2.**
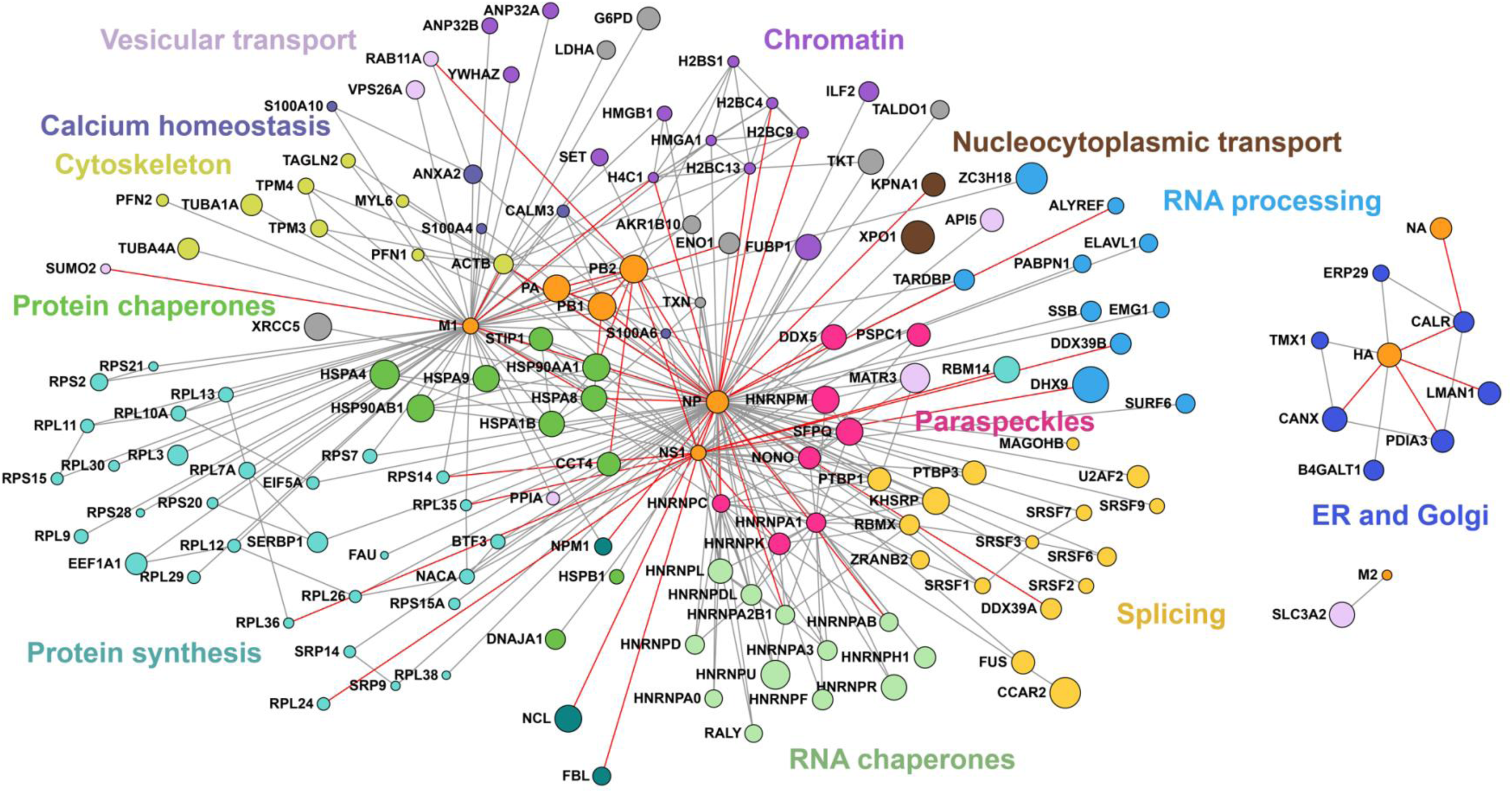
Cross-linking network derived at an FDR threshold of 2% at both residue- and protein-pair levels with additional information mapped. **a,** Cross-linking network highlighting known PPIs with red lines. Host proteins not linked to viral proteins are not shown for clarity. Viral proteins are shown in orange, host proteins are coloured according to the functional category as in Figure 1.

**Extended Data Fig. 3.**
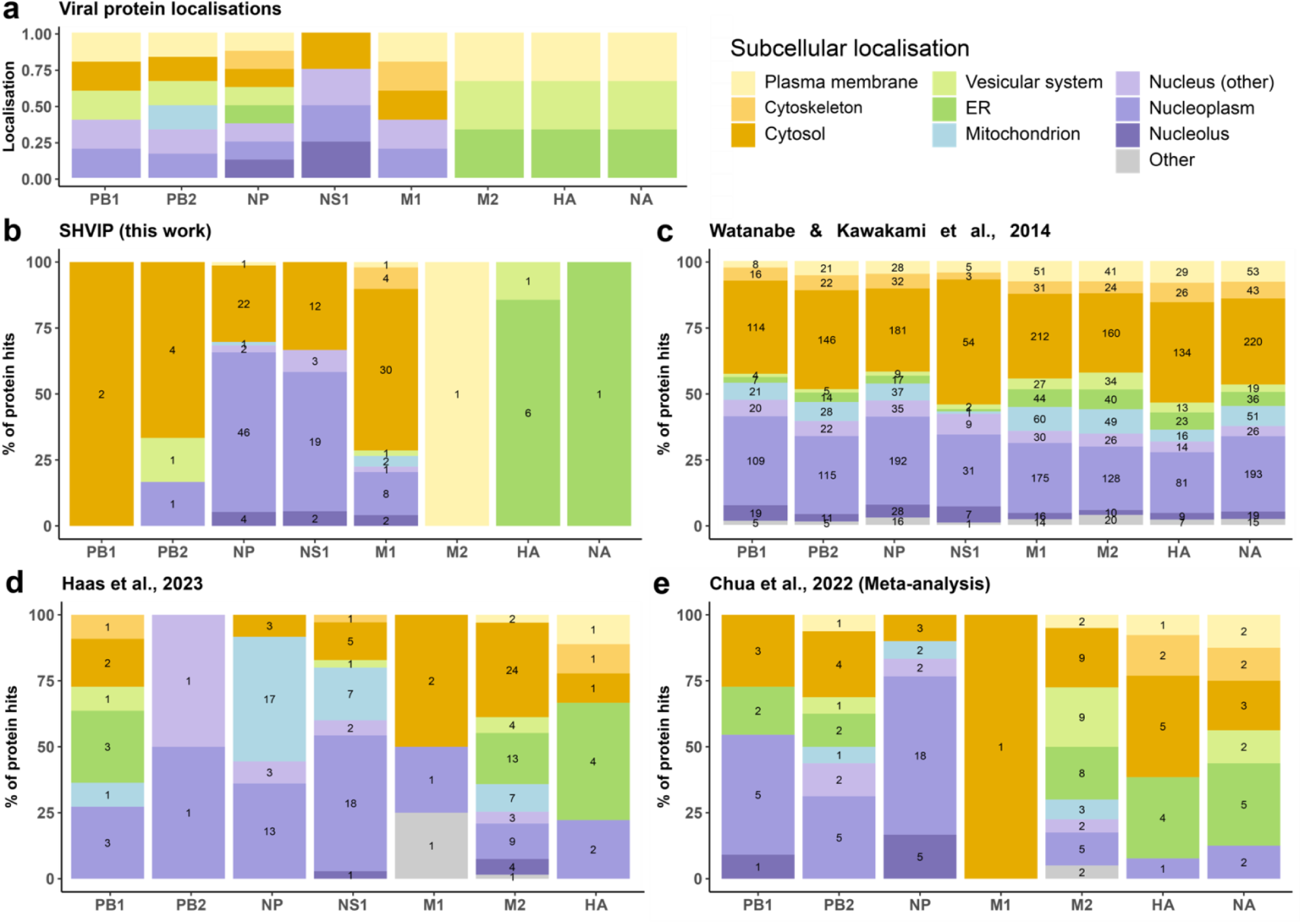
Comparative analysis of subcellular localisations of host interactors of IAV proteins across studies. **a,** Reported subcellular localisations of viral proteins, curated manually from published literature^2,16,38,41,42,51,55,56,137–145^ and databases (UniProt). **b**, SHVIP (this work): In-cell crosslinking during IAV infection captures endogenous host–viral interactions with spatial resolution. **c**, Watanabe & Kawakami et al., 2014: Individual FLAG-tagged viral proteins of A/WSN/33 (H1N1) were overexpressed in HEK293 cells and subjected to anti-FLAG immunoprecipitation followed by mass spectrometry^19^. **d**, Haas et al., 2023: A large-scale AP-MS study using codon-optimised, 2X-Strep-tagged viral proteins expressed in three cell types (A549, NHBE, THP-1) across pH1N1, H3N2 and H5N1 strains^21^. **e**, Chua et al., 2022: A meta-analysis combining host interactors reported across AP-MS, Y2H, RNA-IP, and predictive studies ^40^. Host proteins were considered only if reported in at least three datasets. Only viral proteins identified in SHVIP are shown for other datasets. Bar plots show the percentage distribution of host protein hits (potential interactors) by subcellular compartment for each viral protein. Protein hit counts (n) are overlaid on each bar segment.

**Extended Data Fig. 4.**
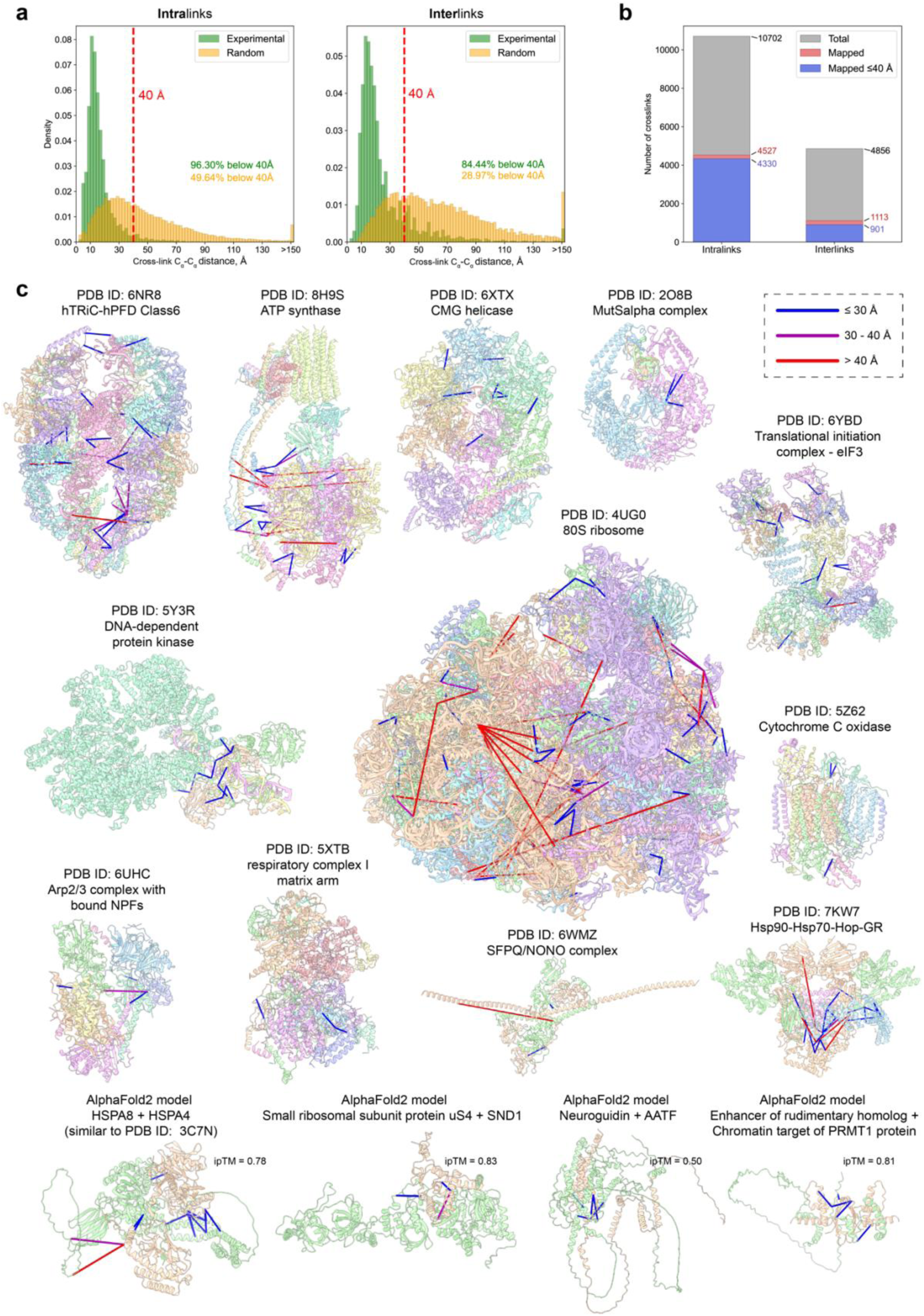
Validation of XL-MS data by available 3D structures and AlphaFold models of cross-linked host PPIs. **a,** Distance histogram of cross-link lengths mapped to 3D structures and AlphaFold. Random distribution corresponds to a random sample of all lysine-residue pairs. The red dashed line indicates the 40 Å threshold. **b,** Bar diagram showing the total number of inter- and intra-cross-links (gray), how many of them could be successfully mapped to PDB structures (red) and how many satisfy the Cα-Cα. **c,** Satisfied cross-links (Cα-Cα distance lower than 30 Å) are coloured blue, cross-links in the range from 30 Å to 40 Å are coloured magenta, while cross-links longer than that are coloured red. **d,** Example AlphaFold2-multimer models with cross-links mapped and coloured as in **(c).**

**Extended data Fig. 5.**
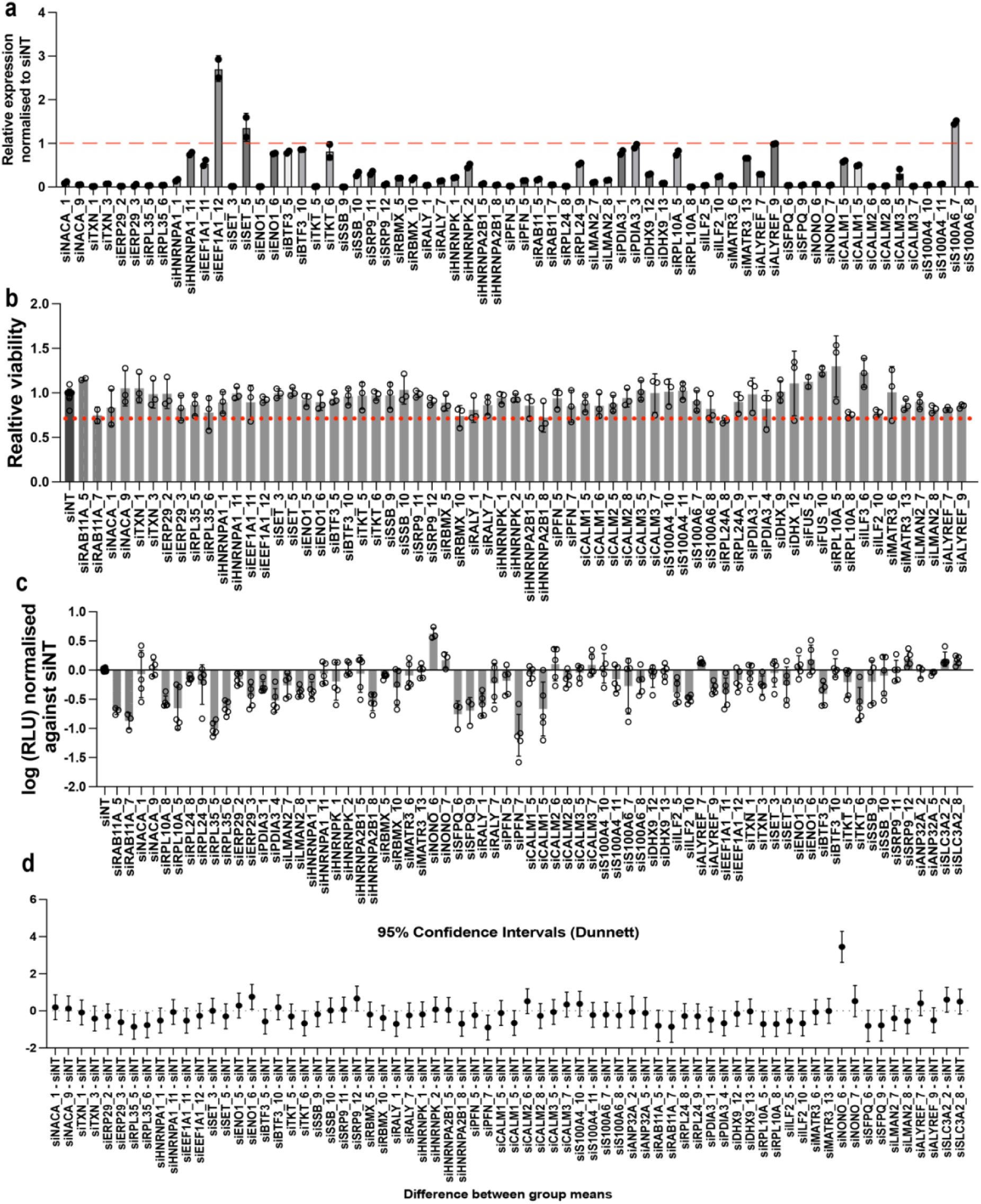
The results of the siRNA screen. **a,** Knockdown efficiency of target genes in A549 cells using siRNA. Efficiency was assessed by qPCR. Data is represented as relative expression of target gene normalised to the non-targeting control (siNT) (n=3). The dashed red line indicates the value obtained for the control. **b,** Viability was measured using the CellTiter-Glo assay and normalised to the non-targeting control (n = 3). The red dashed line indicates a 70% viability cut-off; siRNAs below this threshold were excluded from further analysis. **c,** Luciferase assay in A549 cells with target gene knockdown, infected with recombinant WSN (MOI 0.01, PB2-T2A-NanoLuc) for 48 h. Data are normalised to siNT (n = 4). Error bars represent mean ± SD, with significance determined by ANOVA (Dunnett’s test). **d,** Confidence interval (CI) plot for luciferase activity showing comparisons between knockdowns and siNT. CIs were calculated using ANOVA (Dunnett’s test, n = 4). Data are presented as mean ± 95% CI.

**Extended Data Fig. 6.**
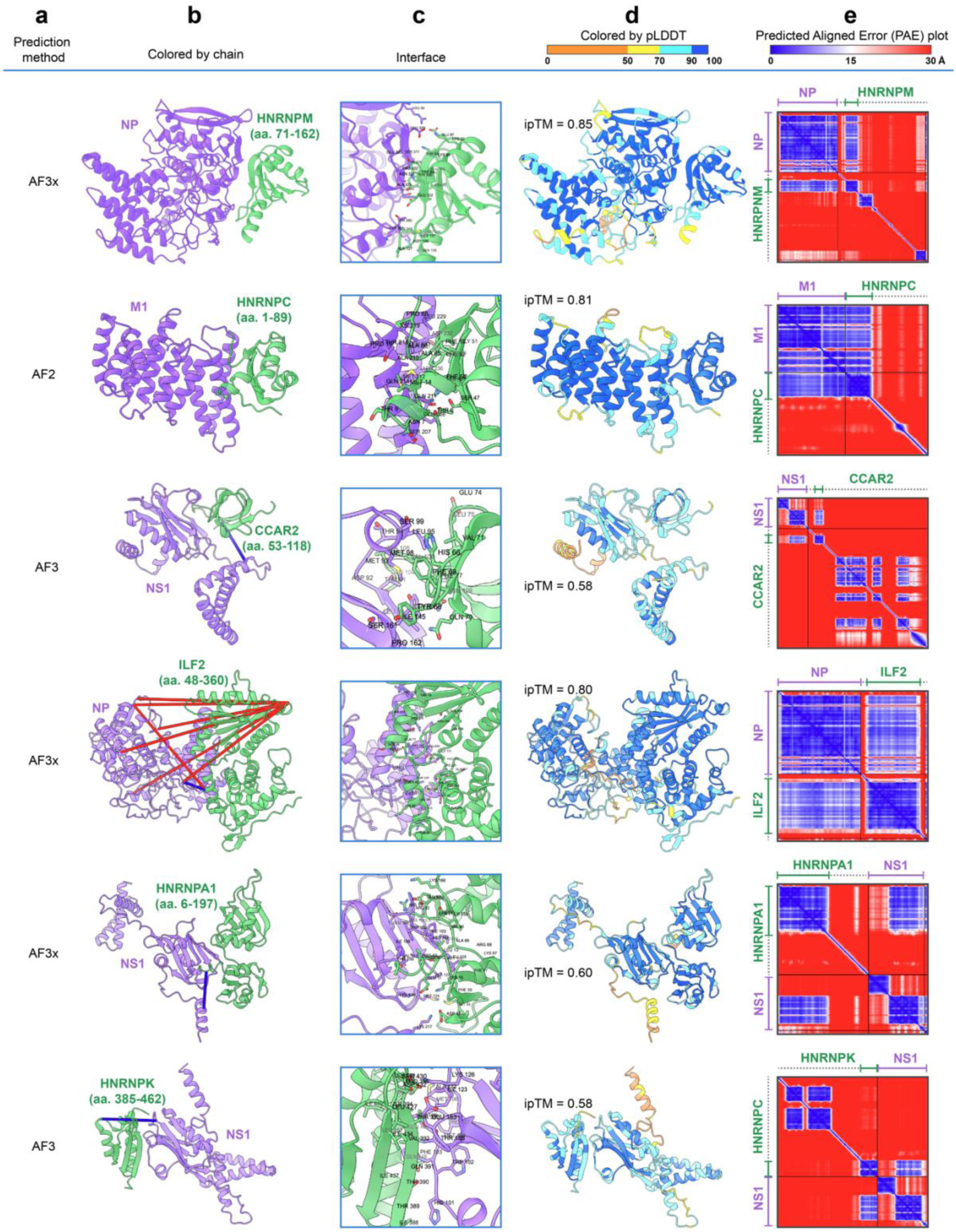
Structural models of IAV proteins in complex with human proteins. Each row presents the structural model of one of the following protein pairs: NP - HNRNPM, M1 - HNRNPC, NS1 - CCAR2, NP - ILF2, NS1 - HNRNPA1, and NS1 - HNRNPK. **a,** The method used for model generation: AlphaFold2-Multimer (AF2), AlphaFold3 (AF3), and AF3x. **b,** Models, where the purple chain corresponds to the IAV protein and the green chain to the human protein. For human proteins, only fragments that confidently interact with the pathogen protein are included. The range of visible amino acids (aa.) is annotated as the first to the last residue shown. Identified interchain cross-links, if present for the shown protein fragments, are marked with lines. Cross-links satisfying the Cα-Cα distance threshold (<30 Å) are coloured blue, while longer cross-links are coloured red. **c,** Close up views on interaction sites, with residues shown that have van der Waals radius intersections greater than 0.4 Å with residues from another chain. **d,** Models coloured according to the predicted local distance difference test (pLDDT), reflecting confidence in the predicted residues (higher values indicate greater confidence). The interface predicted template modelling (ipTM) score is also indicated. **e,** Predicted aligned error (PAE) heat maps, where both axes represent residues from the N-terminus to the C-terminus (left to right and top to bottom). The colour of each matrix position represents the confidence in the spatial arrangement of residues relative to one another, with lower errors (blue) indicating higher confidence. Axes parts corresponding to the model chains are coloured to match the chain colouring in the second column. Residues of the host protein hidden from view in panels **b** and **d** are indicated by dashed lines.

**Extended Data Fig. 7.**
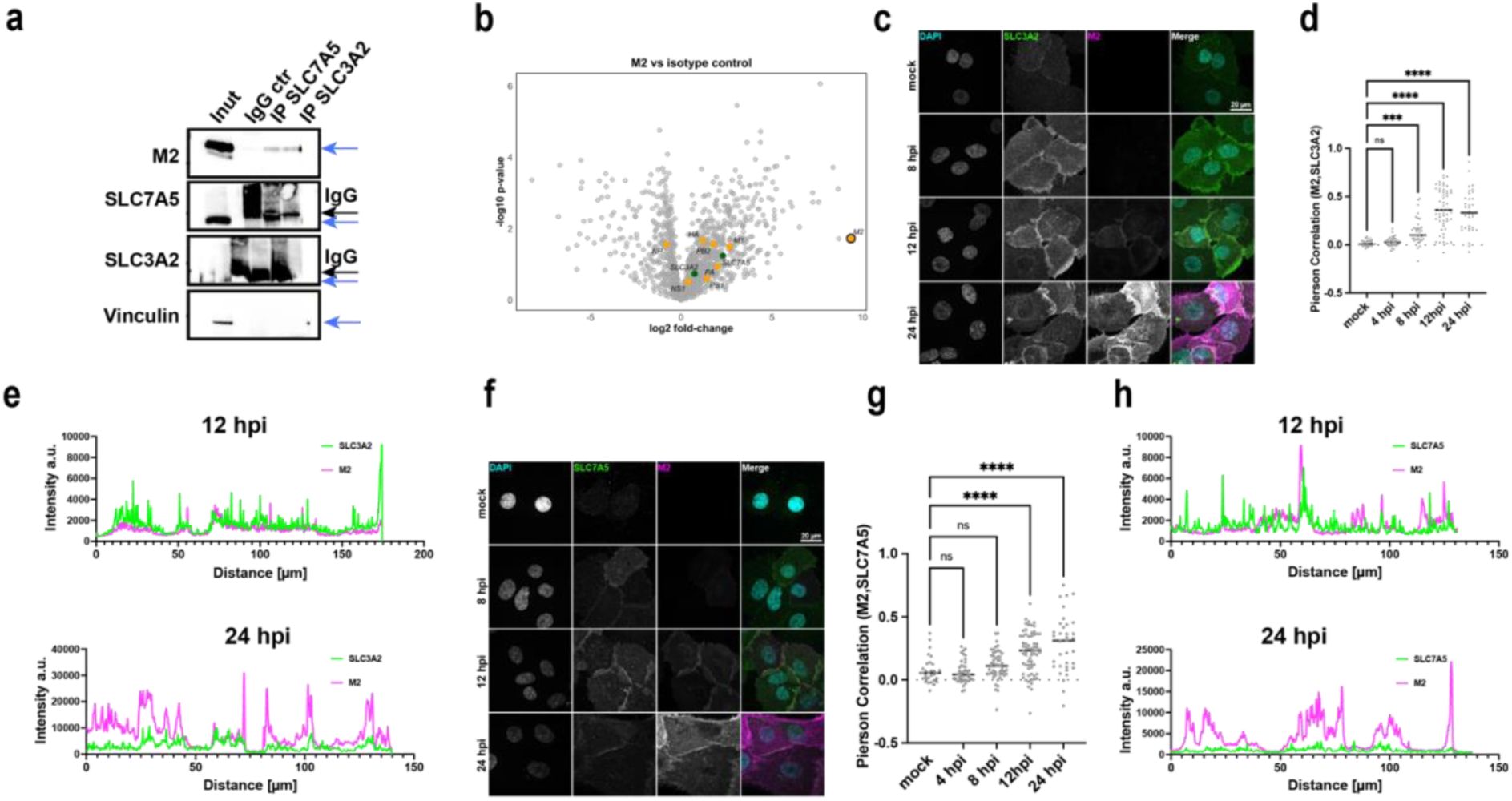
LAT1 in IAV infection. **a,** IP of SLC7A5 and SLC3A2 in A549 cells infected with WSN (MOI 3, 14 hpi) using antibodies against SLC7A5 and SLC3A2 or an isotype control (IgG ctr). Immunoprecipitants were analysed by Western blot for M2, SLC7A5, SLC3A2, and vinculin (negative control). Blue arrows indicate protein of interest, black ones – IgG. **b,** Volcano plots showing fold-change (log2) versus significance (-log10 p-value) (n=3) for M2 versus isotype control. Viral proteins are depicted in orange, SLC7A5 and SLC3A2 – in green. **c,** Maximum intensity projection of primary HBEpCs cells showing colocalisation of SLC3A2 (green) and M2 (magenta). Images were taken at different timepoints during infection and show the distribution of SLC3A2 and M2 in the cells. The maximum intensity projection was generated from z-stacks to visualise overall protein localisation. **d,** Pearson correlation coefficient between SLC3A2 and M2 in both the membrane and cytoplasm of HBEpCs cells. Correlation was calculated to assess the degree of colocalisation between these two proteins using Cell Profiler with 15 % threshold. Statistical analysis was performed using a Kruskal–Wallis test (P < 0.0001) followed by Dunn’s multiple comparisons. Significant increases were observed at 8, 12, and 24 hpi compared to mock (***P = 0.0002, ****P < 0.0001). No significant difference was observed at 4 hpi. n = 31–54 cells per condition. **e**, Intensity profile of a representative plane showing the localisation of SLC3A2 and M2 in several HBEpCs cells at 12 and 24 hpi. The profile represents the fluorescence intensity across a cross-section of the cells. **f,** Maximum intensity projection of HBEpCs cells showing colocalisation of SLC7A5 (green) and M2 (magenta). Images were taken at different timepoints during infection and show the distribution of SLC7A5 and M2 in the cells. The maximum intensity projection was generated from z-stacks to visualise overall protein localisation. **g,** Pearson correlation coefficient between SLC7A5 and M2 in both the membrane and cytoplasm of HBEpCs cells. Correlation was calculated to assess the degree of colocalisation between these two proteins using Cell Profiler with 15 % threshold. Statistical analysis was performed using a Kruskal–Wallis test (P < 0.0001) followed by Dunn’s multiple comparisons. Significant increases in colocalisation were observed at 12 and 24 hpi compared to mock (****P < 0.0001), while changes at 4 and 8 hpi were not significant. n = 31–68 cells per condition. **h,** Intensity profile of a representative plane showing the localisation of SLC7A5 and M2 in several HBEpCs cells at 12 and 24 hpi. The profile represents the fluorescence intensity across a cross-section of the cells.

**Extended Data Fig. 8.**
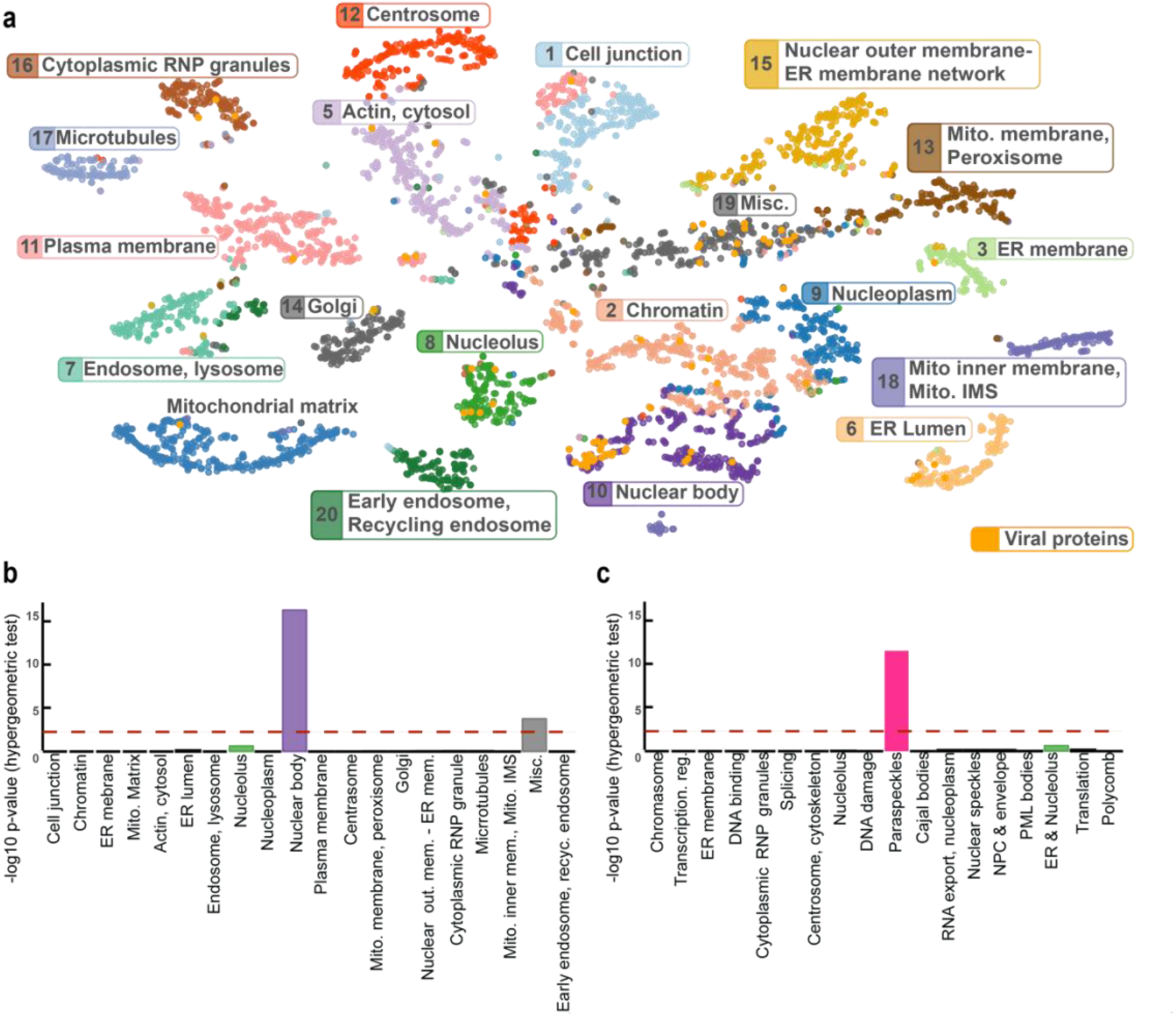
Spatial proteomic context of IAV–host protein interactions. **a,** Mapping of host proteins crosslinked to IAV proteins onto the proximity-dependent biotinylation map of the human proteome^76^. Each coloured cluster represents a spatially resolved cellular compartment. Crosslinked host proteins were overlaid onto the map to infer their spatial context. **b,** Enrichment analysis of IAV-crosslinked host proteins using the Go et al. proximity labelling organelle map^76^. Significant enrichment was observed in the nuclear bodies (purple bar). **c,** Spatial enrichment using the updated nuclear body proteome dataset^77^, revealing a strong enrichment in paraspeckles.

**Extended Data Fig. 9.**
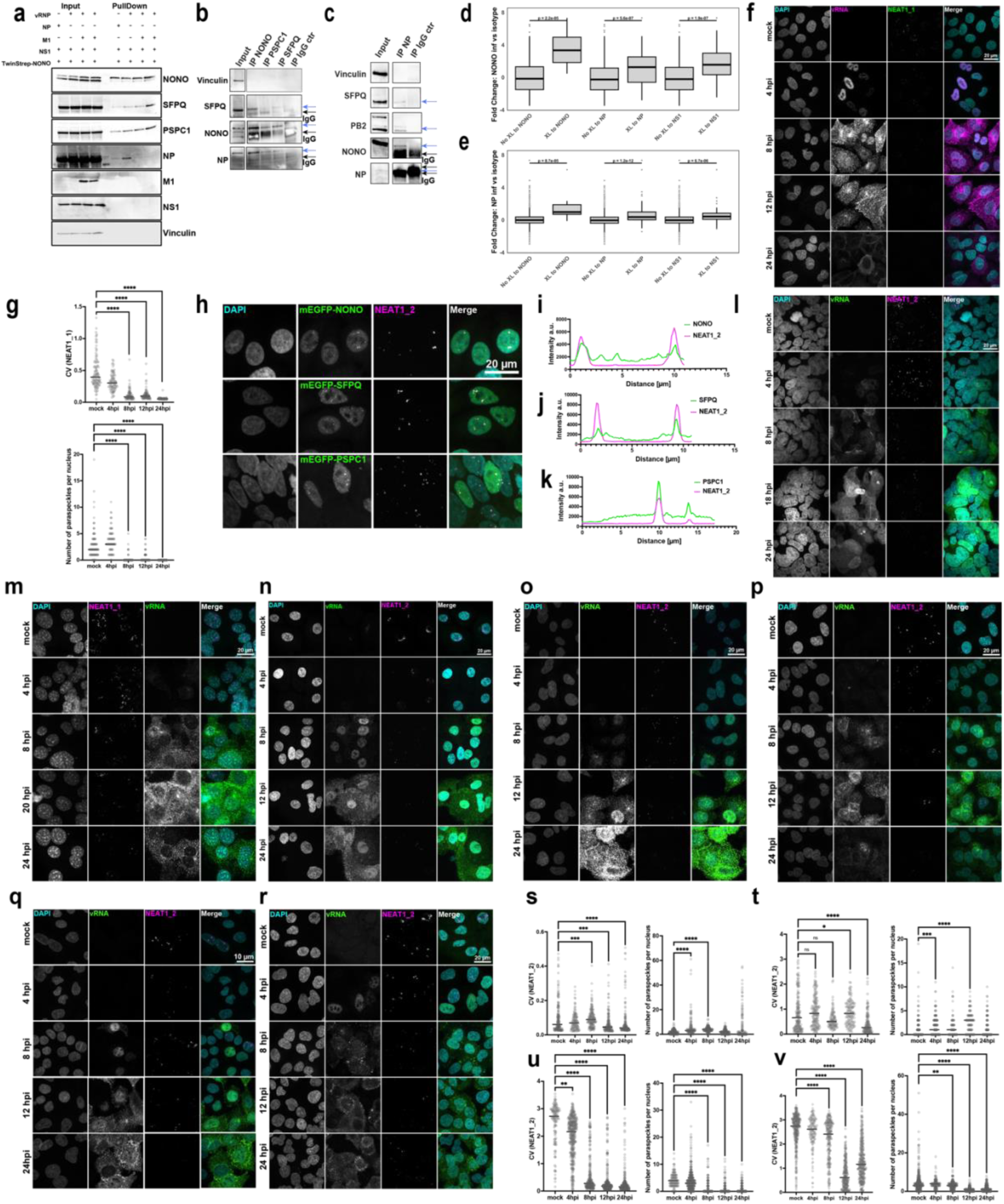
Analysis of paraspeckles during IAV infection. **a,** TwinStrep-tagged NONO was overexpressed in HEK 293T cells alongside the viral proteins NP, M1, and NS1, or the vRNP complex for 24 hours. Streptactin beads were used for pull-down to isolate the TwinStrep-NONO complex. Input and pull-down samples were analysed by western blot. Antibodies against NONO, SFPQ, and PSPC1 were used to confirm successful pulldown of paraspeckle proteins. Antibodies against viral proteins NP, M1, and NS1 were employed to assess their interaction with the NONO complex. **b,** Immunoprecipitation of paraspeckle proteins NONO, SFPQ and PSPC1 and IgG control in A549 infected cells (WSN, MOI=3, 14 hpi). The proteins of interest are annotated with the blue arrows and the IgG bands with the black ones. Vinculin was used as a negative control. **c**, Immunoprecipitation of NP and IgG control in A549 infected cells (WSN, MOI=3, 14 hpi). The proteins of interest are annotated with the blue arrows and the IgG bands with the black ones. NP levels are not distinguishable from IgG bands, therefore PB2 was used as a positive control for IP. Vinculin was used as a negative control. **d**, AP-MS analysis of NONO vs. isotype in A549 cells (WSN, MOI=3, 14 hpi) control showing fold changes for enrichment of NONO, NP, and NS1 cross-linked partners identified in SHVIP, and those not cross-linked (n=3). Statistical analysis was performed using the Wilcoxon test in R. Significance levels: ***p < 0.001, **p < 0.01, *p < 0.05. **e**, AP-MS analysis of NP vs. isotype in A549 cells (WSN, MOI=3, 14 hpi) control showing fold changes for enrichment of NONO, NP, and NS1 cross-linked partners identified in SHVIP, and those not cross-linked (n=3). Statistical analysis was performed using the Wilcoxon test in R. Significance levels: ***p < 0.001, **p < 0.01, *p < 0.05. **f,** Representative maximum projection of confocal images showing NEAT1_1 (green), vRNA (magenta), and DAPI (cyan) staining in mock and infected A549 cells at 4, 8, 12, and 24 hpi. Scale bar = 20 μm. **g,** CV of NEAT1_1 signal intensity and the number of paraspeckles per nucleus across different time points post-infection. n = 91–114 cells per condition. **h,** Maximum projection images of confocal images of lentiviral cell lines expressing mEGFP-tagged SFPQ, NONO, or PSPC1, and their colocalisation with NEAT1_2 (magenta). Scale bar = 20 μm. **i-k,** Line profiles of colocalisation: Gray value intensity profiles of NEAT1_2 (magenta) and mEGFP-tagged proteins (green) showing colocalisation across distance (in microns): (**i**) NONO, (**j**) SFPQ, and (**k**) PSPC1. **(l-n)** Representative maximum projection of confocal images showing NEAT1_2 (magenta), vRNA (green), and DAPI (cyan) staining in mock and infected Calu-3 (**l**), MEFs (**m**), HEBpC (**n**) cells infected with WSN at 4, 8, 18, and 24 hpi. Scale bar = 20 μm. **(o-p)** Representative maximum projection of confocal images showing NEAT1_2 (magenta), vRNA (green), and DAPI (cyan) staining in mock and infected A549 (**o**), HEBpC (**p**) cells infected with pdm09 at 4, 8, 18, and 24 hpi. Scale bar = 20 μm. **(q-r)** Representative maximum projection of confocal images showing NEAT1_2 (magenta), vRNA (green), and DAPI (cyan) staining in mock and infected A549 (q), HBEpC (r) cells infected with H3N2/Aichi at 4, 8, 18, and 24 hpi. Scale bar = 20 μm. **(s)** CV of NEAT1_2 signal intensity and the number of paraspeckles per nucleus in A549 cells during pdm09 infection at various time points. n = 136–196 cells per condition. **(t)** CV of NEAT1_2 signal intensity and the number of paraspeckles per nucleus in primary HBEpC cells during pdm09 infection at various time points. n = 134–393 cells per condition. **(u)** CV of NEAT1_2 signal intensity and the number of paraspeckles per nucleus in A549 cells during H3N2/Aichi infection at various time points. n = 98–167 cells per condition. **(v)** CV of NEAT1_2 signal intensity and the number of paraspeckles per nucleus in HBEpC cells during H3N2/Aichi infection at various time points. n = 148–379 cells per condition. Data from panels (**g, s-v**) are shown as individual values. Statistical analysis was performed using one-way ANOVA with Dunnett’s multiple comparisons test, two-sided. Significance levels: *p < 0.05, **p < 0.01, ***p < 0.001, \*\*\**p < 0.0001*.

**Extended Data Fig. 10.**
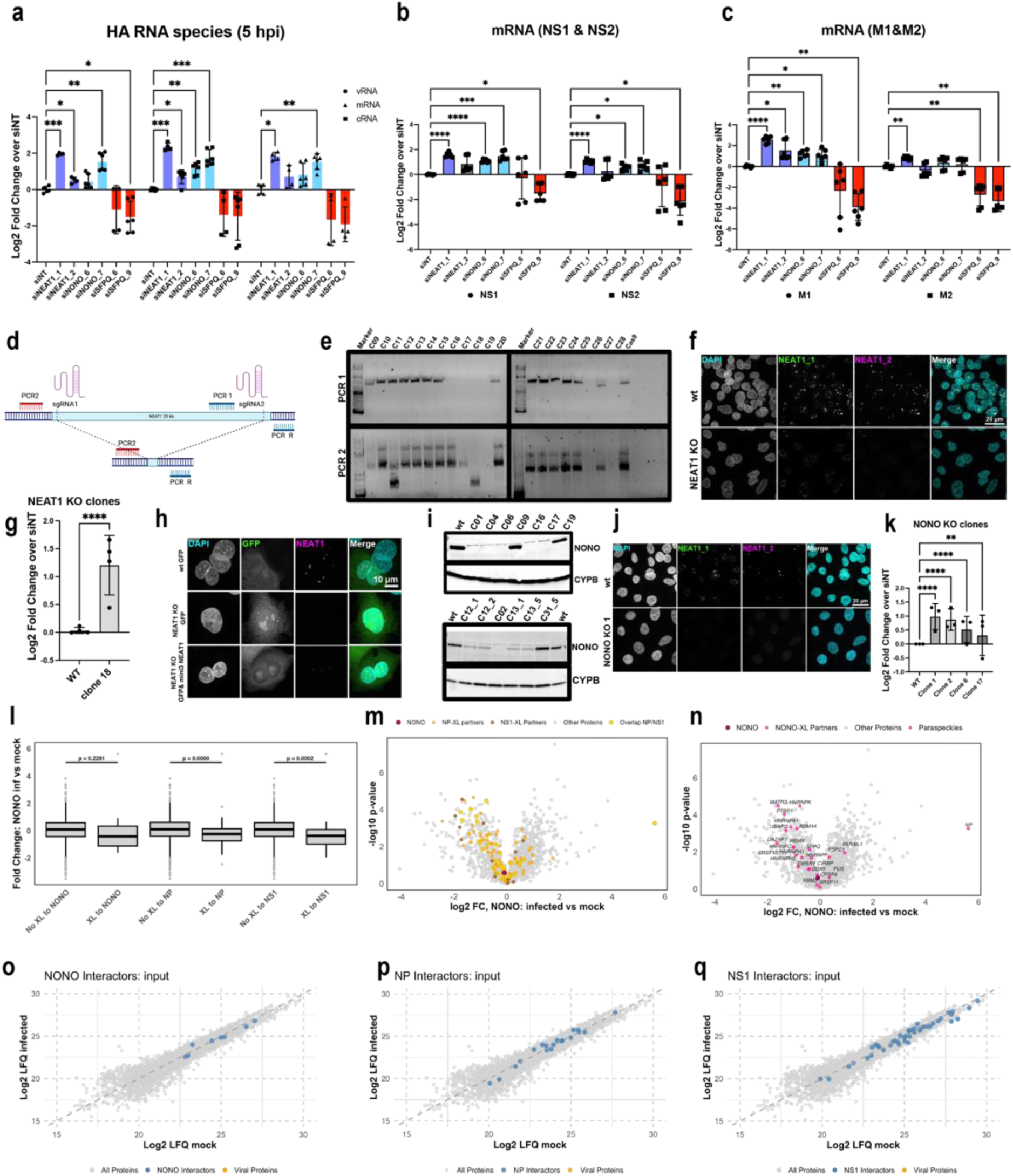
Role of NEAT1 and NONO during IAV infection. **a,** qPCR analysis of viral RNA species of HA fragment normalised to GAPDH in A549 cells with knockdowns of paraspeckle proteins (WSN, MOI 3, 6hpi, n=3). **b-c,** qPCR analysis of spliced and unspliced viral RNA species in A549 cells with knockdowns of paraspeckle proteins (WSN, MOI 3, 6hpi, n=3): NS, NS2 (**b**) and M1, M2 (**c**). **d,** Strategy for NEAT1 KO Generation: Schematic of the CRISPR/Cas9-based approach to generate NEAT1 KO in A549 cells. Two sgRNAs are used to target NEAT1. PCR1 amplifies the region only in wild type (WT) or heterozygous clones, whereas PCR2 detects the shorter product (approximately 500 bp) in KO clones. For heterozygous clones, both the shorter KO product and the long (23 kb) WT product may appear. *Created in BioRender.* https://BioRender.com/j78u936. **e,** PCR Screening of Clones: Agarose gel showing PCR1 and PCR2 results for various clones. Clone 18 exhibits only the shorter PCR2 product, indicating a homozygous NEAT1 KO, while Clone 11 shows both products, indicating heterozygosity. **f,** Maximum projection of confocal images of NEAT1: Representative images showing the localisation of NEAT1_1 (green) and NEAT1_2 (magenta) in WT and NEAT1 KO (Clone 28) cells. DAPI (cyan) counterstain nuclei. Scale bar = 20 μm. **g,** Quantification of luciferase activity in WT and NEAT1 KO cells. **h,** Overexpression of miniNEAT1 3 in A549 NEAT1 KO cells using electroporation partially rescued paraspeckle formation. **I,** Western blot showing NONO protein levels in WT and NONO KO clones generated using sgRNA1 (above) sgRNA2 (below). CYPB serves as a loading control. **j**, Maximum projection of confocal images of NEAT1: IF images of NEAT1_1 (green) and NEAT1_2 (magenta) in WT and NONO KO (Clone 2) cells. DAPI (cyan) counterstains nuclei. Scale bar = 20 μm. **k,** Quantification of luciferase activity in WT and NONO KO A549 cells. **l,** Fold-change in NONO AP_MS (infected vs. mock) identified in AP-MS, including NONO, NP, and NS1 cross-linked partners. Data represent mean ± SD (n = 3). Statistical significance was determined using a Wilcoxon test. **m,** Volcano plot showing fold-change (log2 FC) vs. statistical significance (-log10 p-value) of NONO AP-MS in infected vs. mock samples. Paraspeckle proteins and NONO cross-linked partners are highlighted in pink. **n,** Volcano plot showing fold-change (log2 FC) vs. statistical significance (-log10 p-value) of NONO AP-MS in infected vs. mock samples. NP-XL, NS1-XL, and overlapping interactors are colour-coded. **o-q,** Scatter plots of Log2 LFQ values comparing infected and mock input samples in AP-MS (NONO inf and NONO mock). (o) NONO XL-partners, (p) NP XL-partners, and (q) NS1 XL-partners are highlighted in blue. Data from panels (**a-c,g,k**) represent mean ± SD (n = 3). Statistical analysis was performed using mixed effect model (a-c) the one-way ANNOVA (g,k) with Dunnett multiple comparison adjustment, two-sided. Significance levels: ***p < 0.001, **p < 0.01, *p < 0.05, ns = not significant.

**Extended Data Table 1 (Excel file) Cross-linking data set. a,** Cross-links filtered at 2% residue-pair FDR level. **b,** Protein pairs filtered at 2% protein-pair FDR level. **c,** Cross-linking data filtered at both 2% residue-pair and protein-pair FDR level. **d,** Virus-centric network containing cross-links to viral proteins and between host proteins that cross-linked to viral proteins. **e,** Cross-links filtered at 5% residue-pair FDR level. **f,** Protein pairs filtered at 5% protein-pair FDR level. **g,** Cross-linking data filtered at both 5% residue-pair and protein-pair FDR level.

**Extended Data Table 2 (Excel file) Protein pairs identified as PPIs in previous focused IAV studies.**

**Extended Data Table 3 (Excel file) Functional enrichment analysis. a,** Host proteins cross-linked to viral proteins and used for the enrichment analysis. **b,** Enriched Reactome Pathways. **c,** Enriched GO biological processes. **d,** Enriched GO cellular components. **e,** Enriched GO molecular functions. **f,** Proteins simultaneously enriched among the host proteins in the NONO AP-MS under mock conditions and NP AP-MS infected. See also Figure 7H. **g,** GO biological processes simultaneously enriched for proteins in **(f).** See also Figure 7I.

**Extended Data Table 4 (Excel file) Scores of structural models of IAV-human proteins pairs modelled using AlphaFold. a,** Models built using AlphaFold 3. **b,** Models built using AF3x and all cross-links. **c,** Models built using AF3x and one cross-link at a time. d, Models built using AlphaFold 2. **e,** Selected models built using AlphaFold 3 run 1,000 times with different random generator seeds. **f,** Selected models built using AF3x, one cross-link, and run 1,000 times with different random generator seeds.

**Extended Data Video 1 NONO or SFPQ cellular localisation in A549 cells stably overexpressing mEGFP-NONO or mEGFP-SFPQ infected with WSN at MOI 3 or mock-infected and imaged every 40 min**

