## Supplementary Figures for "Snapshot of in-cell protein contact sites reveals new host factors and hijacking of paraspeckles during influenza A virus infection"

### Supplemental Information.

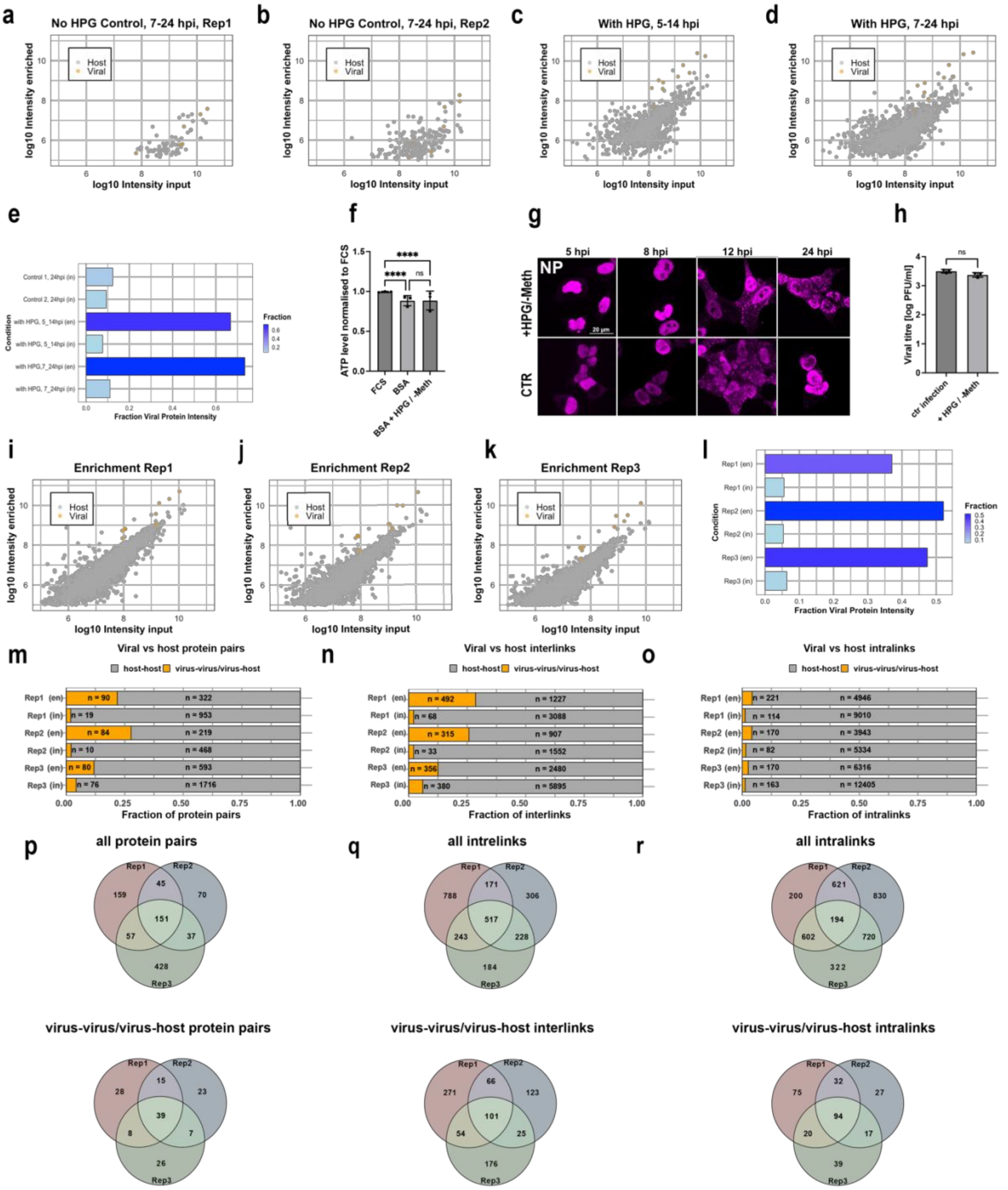

**Extended Data Fig. 1 | Optimisation of SHVIP and enrichment of IAV proteome in the SHVIP experiment. a-d,** Determination of optimal time frames for selective enrichment of viral proteins. Scatter plots represent the correlation between input (x-axis) and enriched (y-axis) log10 iBAQ-values (intensity-based absolute quantification of proteins) of host (grey) and viral (orange) proteins at various time points and conditions. Conditions include no-HPG controls (**a-b**) at 7–24 hpi and HPG-containing samples at 5–14 hpi (**c**) and 7–24 hpi (**d**). L-HPG stands for L-homopropargylglycine. **e,** Fraction of viral protein intensities in total proteomic data under various conditions summarising the data show in **(a-d)**. Bars represent the proportion of viral proteins enriched relative to input proteins across different experimental setups. Enriched samples indicated as en, and input as in. **f,** Viability of A549 cells after incubation in three different media conditions: complete media (FCS), infectious media (BSA), and SHVIP media (BSA L-HPG). Cell viability is normalised to the complete media condition (FCS) (n=3). Data represent mean ± SD. Statistical significance was determined using an ordinary one-way ANOVA. Significance levels are indicated as follows: **** for $p < 0.0001$ and ns for not significant. **g,** Maximum projection of A549 cells infected with WSN at various time points. The viral nucleoprotein (NP) is used as an infection marker (magenta). Cells were treated with either +HPG/-Meth media or standard infectious media (CTR). Meth stands for methionine. **h,** Viral titres at 14 hpi comparing standard infectious media and +HPG/-Meth media protocols. Data represent mean ± SD (n=3). Statistical significance was determined using an ordinary one-way ANOVA. Significance levels are indicated as follows: ns for not significant. **i-k,** Scatter plots represent the correlation between input (x-axis) and enriched (y-axis) log10 iBAQ-values of host (grey) and viral (orange) proteins for three independent replicates. **l,** Fraction of viral protein intensities in total proteomic data in three replicates summarising the data shown in I-K. Bars represent the proportion of viral proteins enriched relative to input proteins across different experimental setups. Enriched samples are indicated as en, and input as in. **m,** Fraction of viral-viral/viral-host and host-host protein pairs across three replicates. **n,** Fraction of viral-viral/viral-host and host-host interlinks across three replicates. **o**, Fraction of viral-viral/viral-host and host-host intralinks across three replicates. **p,** Venn-diagram depicting overlap of all protein pairs (above) and viral-viral/viral host protein pairs among three replicates. **q,** Venn-diagram depicting overlap of all interlinks (above) and viral-viral/viral host interlinks among three replicates. **r,** Venn-diagram depicting overlap of all intralinks (above) and viral-viral/viral host intralinks among three replicates.

**Extended Data Table 1 (Excel file) | Cross-linking data set. a,** Cross-links filtered at 2% residue-pair FDR level. **b,** Protein pairs filtered at 2% protein-pair FDR level. **c,** Cross-linking data filtered at both 2% residue-pair and protein-pair FDR level. **d,** Virus-centric network containing cross-links to viral proteins and between host proteins that cross-linked to viral proteins. **e,** Cross-links filtered at 5% residue-pair FDR level. **f,** Protein pairs filtered at 5% protein-pair FDR level. **g,** Cross-linking data filtered at both 5% residue-pair and protein-pair FDR level.

**Extended Data Table 2 (Excel file) | Protein pairs identified as PPIs in previous focused IAV studies.**

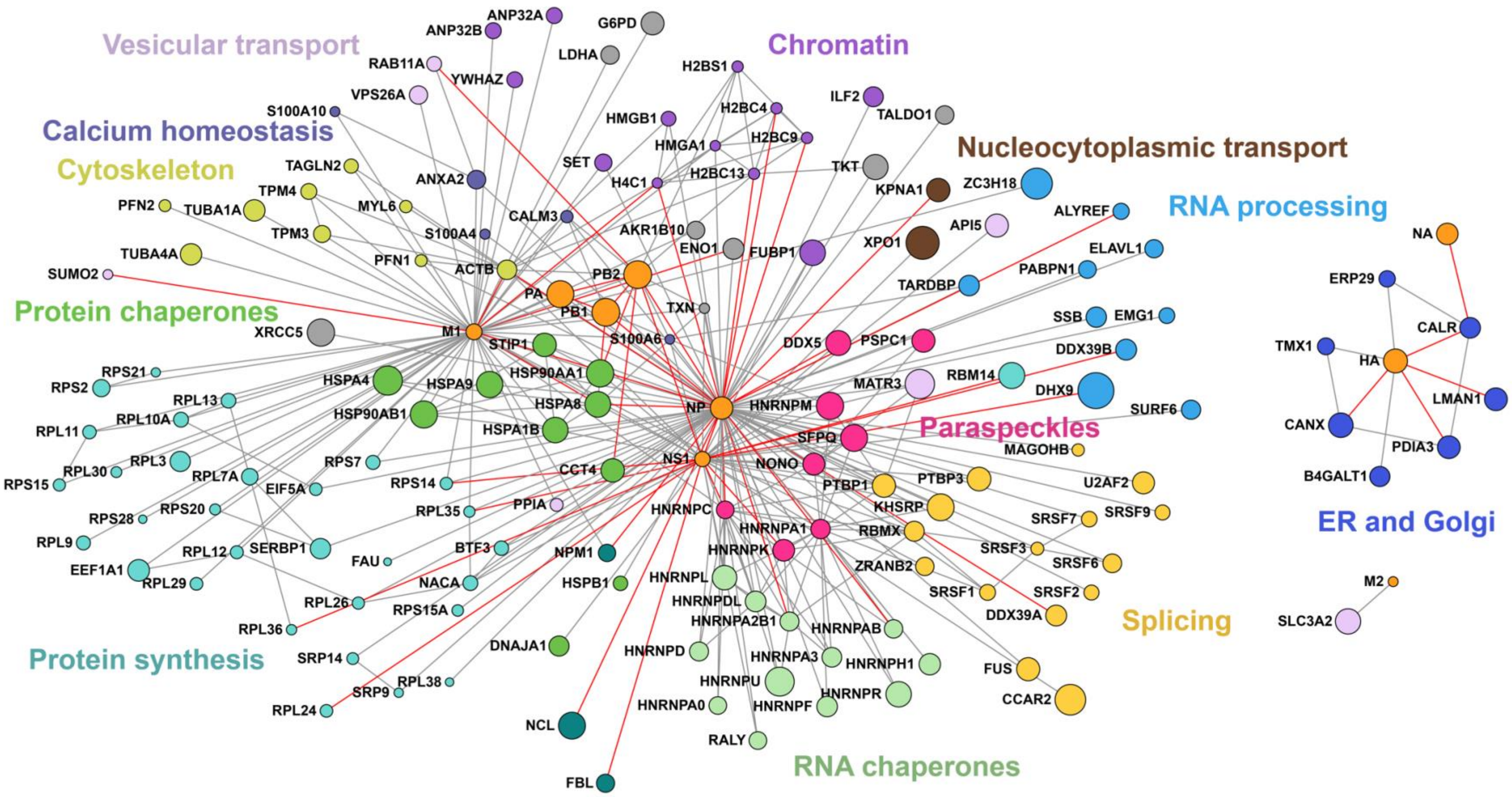


**Extended Data Fig. 2 | Cross-linking network derived at an FDR threshold of 2% at both residue- and protein-pair levels with additional information mapped. a,** Cross-linking network highlighting known PPIs with red lines. Host proteins not linked to viral proteins are not shown for clarity. Viral proteins are shown in orange, host proteins are coloured according to the functional category as in Figure 1.

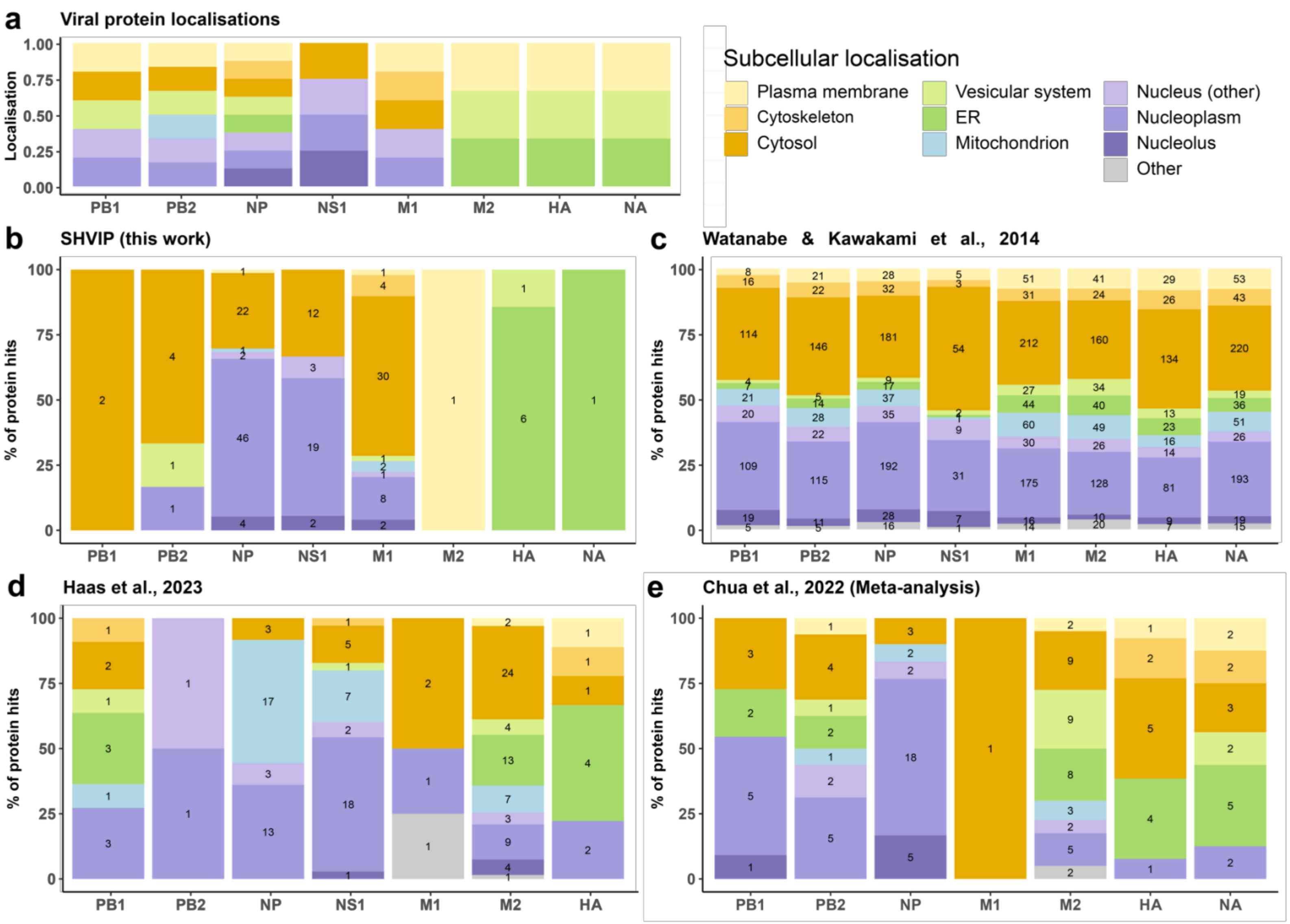
a
Viral protein localisations
Localisation
1.00
0.75
0.50
0.25
0.00
PB1
PB2
NP
NS1
M1
M2
HA
NA
Subcellular localisation
Plasma membrane
Cytoskeleton
Cytosol
Vesicular system
ER
Mitochondrion
Nucleus (other)
Nucleoplasm
Nucleolus
Other
b
SHVIP (this work)
% of protein hits
c
Watanabe & Kawakami et al., 2014
d
Haas et al., 2023
e
Chua et al., 2022 (Meta-analysis)

**Extended Data Fig. 3 | Comparative analysis of subcellular localisations of host interactors of IAV proteins across studies.** **a,** Reported subcellular localisations of viral proteins, curated manually from published literature[2,16,38,41,42,51,55,56,137–145] and databases (UniProt). **b**, SHVIP (this work): In-cell crosslinking during IAV infection captures endogenous host–viral interactions with spatial resolution. **c**, Watanabe & Kawakami et al., 2014: Individual FLAG-tagged viral proteins of A/WSN/33 (H1N1) were overexpressed in HEK293 cells and subjected to anti-FLAG immunoprecipitation followed by mass spectrometry[19]. **d**, Haas et al., 2023: A large-scale AP-MS study using codon-optimised, 2X-Strep-tagged viral proteins expressed in three cell types (A549, NHBE, THP-1) across pH1N1, H3N2 and H5N1 strains[21]. **e**, Chua et al., 2022: A meta-analysis combining host interactors reported across AP-MS, Y2H, RNA-IP, and predictive studies [40]. Host proteins were considered only if reported in at least three datasets. Only viral proteins identified in SHVIP are shown for other datasets. Bar plots show the percentage distribution of host protein hits (potential interactors) by subcellular compartment for each viral protein. Protein hit counts (n) are overlaid on each bar segment.

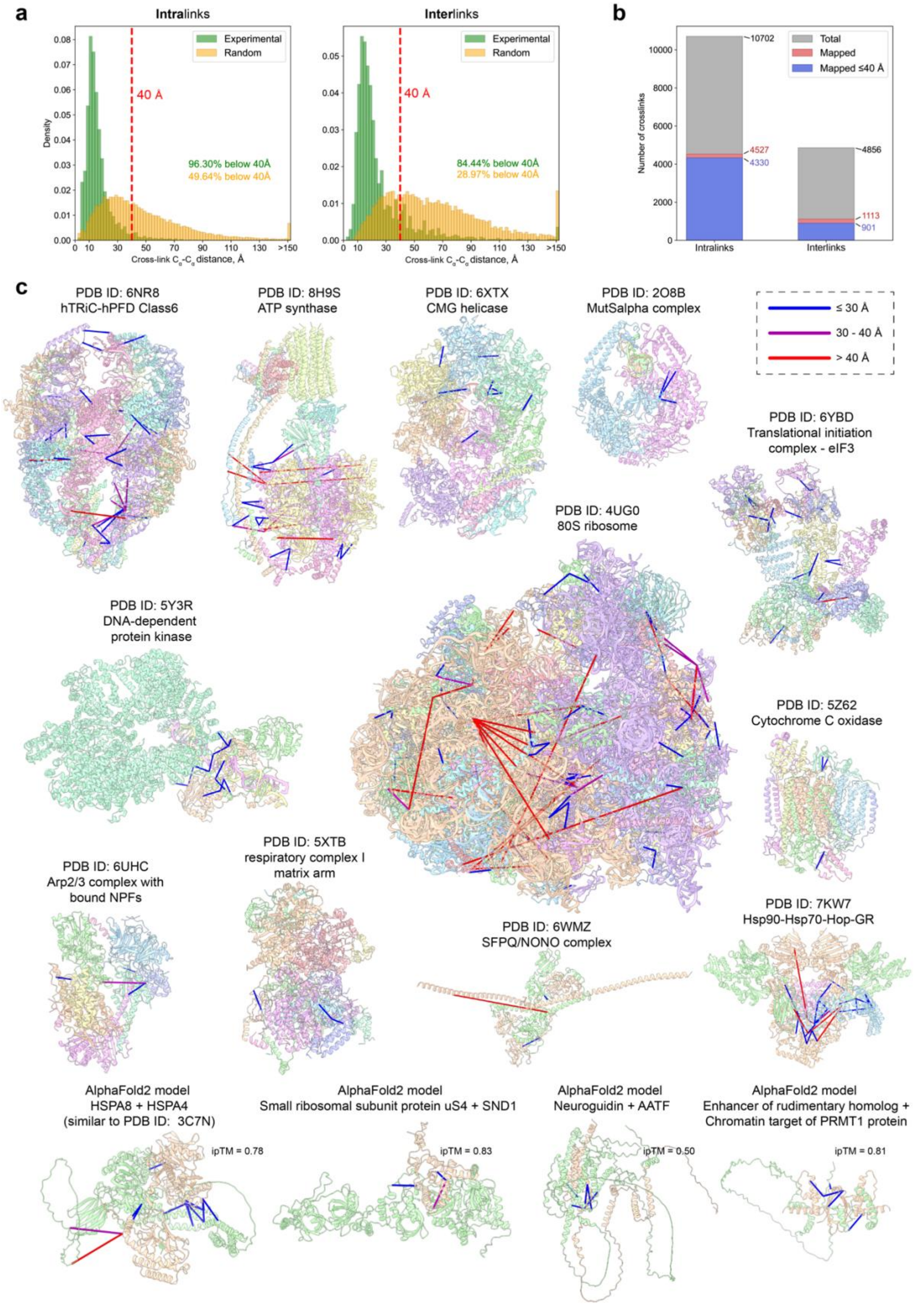


**Extended Data Fig. 4 | Validation of XL-MS data by available 3D structures and AlphaFold models of cross-linked host PPIs. a,** Distance histogram of cross-link lengths mapped to 3D structures and AlphaFold. Random distribution corresponds to a random sample of all lysine-residue pairs. The red dashed line indicates the 40 Å threshold. **b,** Bar diagram showing the total number of inter- and intra-cross-links (gray), how many of them could be successfully mapped to PDB structures (red) and how many satisfy the Cα -Cα. **c,** Satisfied cross-links (Cα-Cα distance lower than 30 Å) are coloured blue, cross-links in the range from 30 Å to 40 Å are coloured magenta, while cross-links longer than that are coloured red. **d,** Example AlphaFold2-multimer models with cross-links mapped and coloured as in **(c).**

**Extended Data Table 3 (Excel file) | Functional enrichment analysis. a,** Host proteins cross-linked to viral proteins and used for the enrichment analysis. **b,** Enriched Reactome Pathways. **c,** Enriched GO biological processes. **d,** Enriched GO cellular components. **e,** Enriched GO molecular functions. **f,** Proteins simultaneously enriched among the host proteins in the NONO AP-MS under mock conditions and NP AP-MS infected. See also Figure 7H. **g,** GO biological processes simultaneously enriched for proteins in **(f).** See also Figure 7I.

**Extended Data Table 4 (Excel file) | Scores of structural models of IAV-human proteins pairs modelled using AlphaFold. a,** Models built using AlphaFold 3. **b,** Models built using AF3x and all cross-links. **c,** Models built using AF3x and one cross-link at a time. d, Models built using AlphaFold 2. **e,** Selected models built using AlphaFold 3 run 1,000 times with different random generator seeds. **f,** Selected models built using AF3x, one cross-link, and run 1,000 times with different random generator seeds.

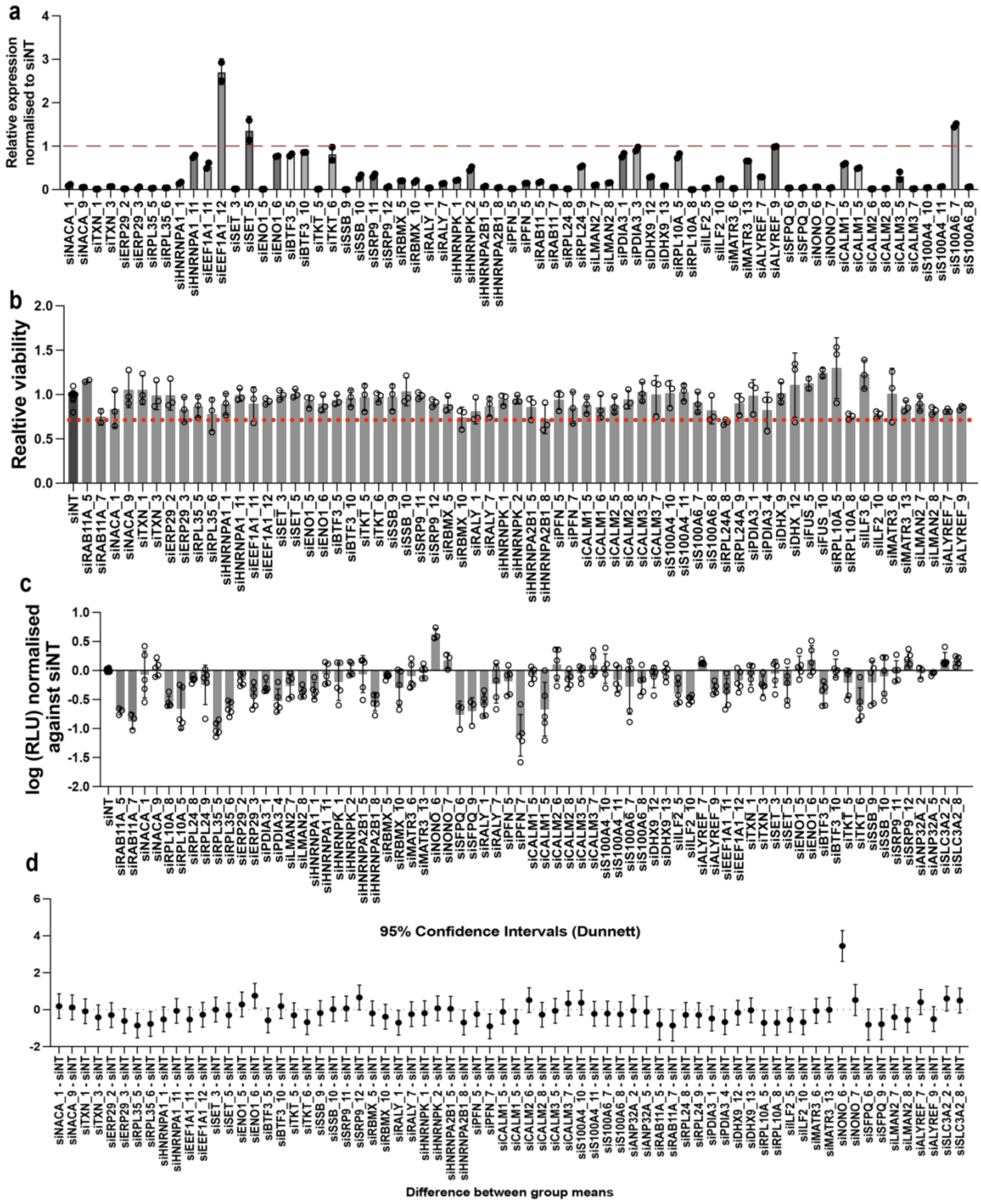


**Extended data Fig. 5 | The results of the siRNA screen. a,** Knockdown efficiency of target genes in A549 cells using siRNA. Efficiency was assessed by qPCR. Data is represented as relative expression of target gene normalised to the non-targeting control (siNT) (n=3). The dashed red line indicates the value obtained for the control. **b,** Viability was measured using the CellTiter-Glo assay and normalised to the non-targeting control (n = 3). The red dashed line indicates a 70% viability cut-off; siRNAs below this threshold were excluded from further analysis. **c,** Luciferase assay in A549 cells with target gene knockdown, infected with recombinant WSN (MOI 0.01, PB2-T2A-NanoLuc) for 48 h. Data are normalised to siNT (n = 4). Error bars represent mean ± SD, with significance determined by ANOVA (Dunnett's test). **d,** Confidence interval (CI) plot for luciferase activity showing comparisons between knockdowns and siNT. CIs were calculated using ANOVA (Dunnett's test, n = 4). Data are presented as mean ± 95% CI.

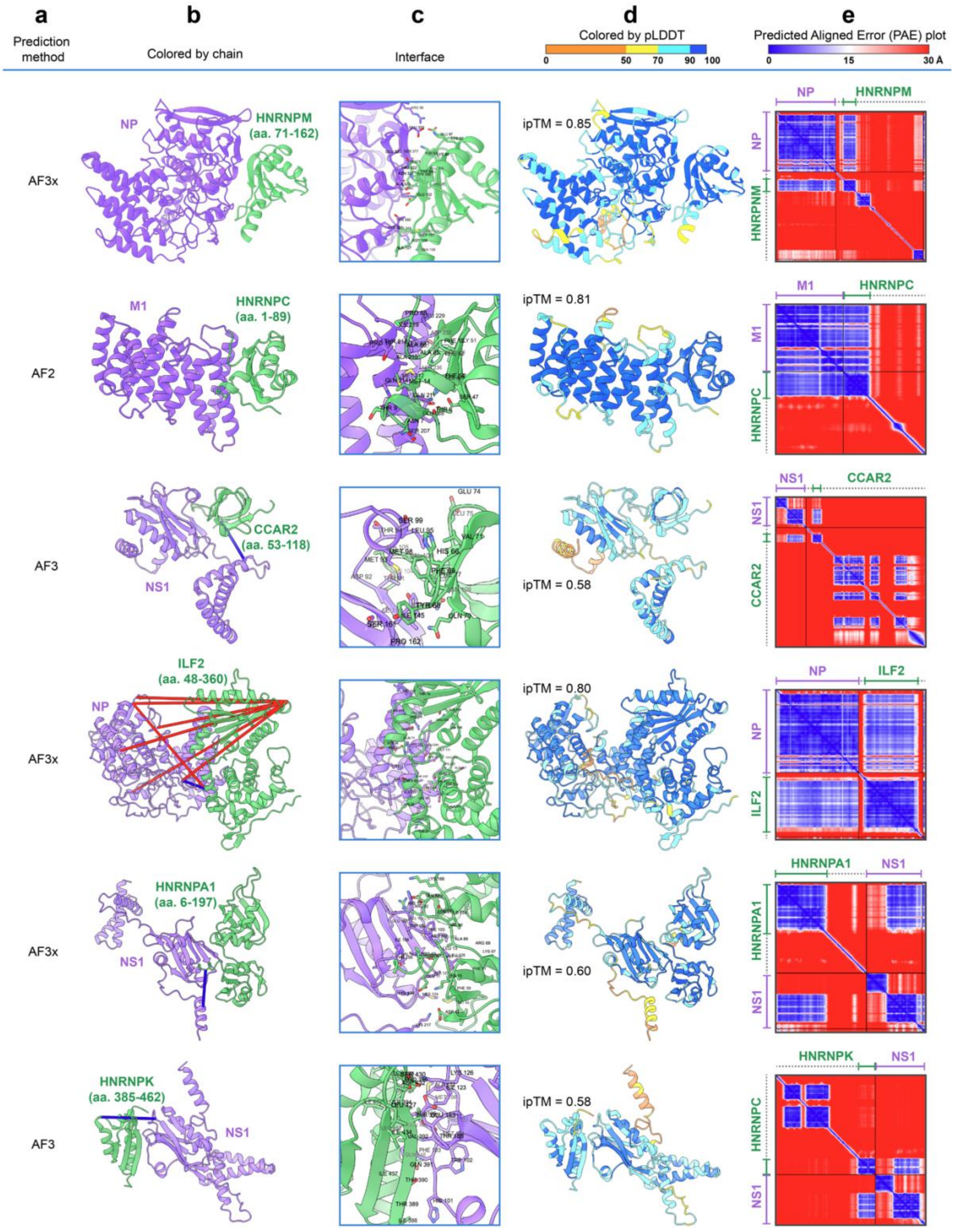


**Extended Data Fig. 6 | Structural models of IAV proteins in complex with human proteins.** Each row presents the structural model of one of the following protein pairs: NP - HNRNPM, M1 - HNRNPC, NS1 - CCAR2, NP - ILF2, NS1 - HNRNPA1, and NS1 - HNRNPK. **a,** The method used for model generation: AlphaFold2-Multimer (AF2), AlphaFold3 (AF3), and AF3x. **b,** Models, where the purple chain corresponds to the IAV protein and the green chain to the human protein. For human proteins, only fragments that confidently interact with the pathogen protein are included. The range of visible amino acids (aa.) is annotated as the first to the last residue shown. Identified interchain cross-links, if present for the shown protein fragments, are marked with lines. Cross-links satisfying the Cα-Cα distance threshold (<30 Å) are coloured blue, while longer cross-links are coloured red. **c,** Close up views on interaction sites, with residues shown that have van der Waals radius intersections greater than 0.4 Å with residues from another chain. **d,** Models coloured according to the predicted local distance difference test (pLDDT), reflecting confidence in the predicted residues (higher values indicate greater confidence). The interface predicted template modelling (ipTM) score is also indicated. **e,** Predicted aligned error (PAE) heat maps, where both axes represent residues from the N-terminus to the C-terminus (left to right and top to bottom). The colour of each matrix position represents the confidence in the spatial arrangement of residues relative to one another, with lower errors (blue) indicating higher confidence. Axes parts corresponding to the model chains are coloured to match the chain colouring in the second column. Residues of the host protein hidden from view in panels **b** and **d** are indicated by dashed lines.

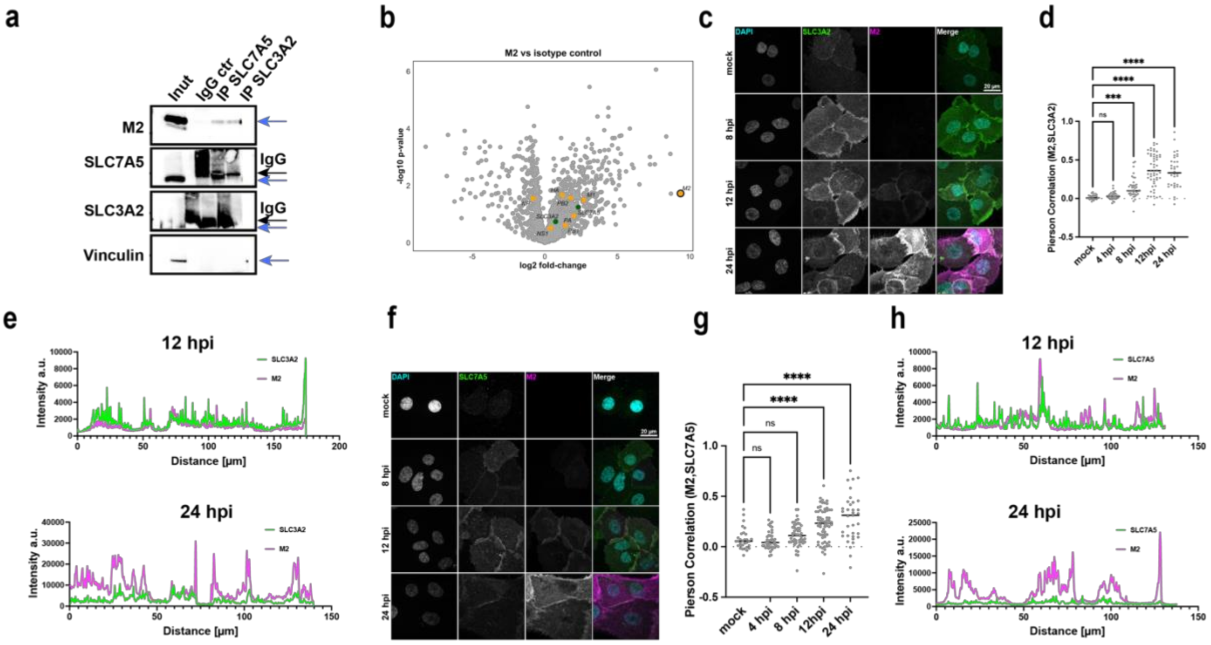


**Extended Data Fig. 7 | LAT1 in IAV infection. a,** IP of SLC7A5 and SLC3A2 in A549 cells infected with WSN (MOI 3, 14 hpi) using antibodies against SLC7A5 and SLC3A2 or an isotype control (IgG ctr). Immunoprecipitants were analysed by Western blot for M2, SLC7A5, SLC3A2, and vinculin (negative control). Blue arrows indicate protein of interest, black ones – IgG. **b,** Volcano plots showing fold-change (log2) versus significance (-log10 p-value) (n=3) for M2 versus isotype control. Viral proteins are depicted in orange, SLC7A5 and SLC3A2 – in green. **c,** Maximum intensity projection of primary HBEpCs cells showing colocalisation of SLC3A2 (green) and M2 (magenta). Images were taken at different timepoints during infection and show the distribution of SLC3A2 and M2 in the cells. The maximum intensity projection was generated from z-stacks to visualise overall protein localisation. **d,** Pearson correlation coefficient between SLC3A2 and M2 in both the membrane and cytoplasm of HBEpCs cells. Correlation was calculated to assess the degree of colocalisation between these two proteins using Cell Profiler with 15 % threshold. Statistical analysis was performed using a Kruskal–Wallis test ($P < 0.0001$) followed by Dunn's multiple comparisons. Significant increases were observed at 8, 12, and 24 hpi compared to mock (***$P = 0.0002$, ****$P < 0.0001$). No significant difference was observed at 4 hpi. n = 31–54 cells per condition. **e**, Intensity profile of a representative plane showing the localisation of SLC3A2 and M2 in several HBEpCs cells at 12 and 24 hpi. The profile represents the fluorescence intensity across a cross-section of the cells. **f,** Maximum intensity projection of HBEpCs cells showing colocalisation of SLC7A5 (green) and M2 (magenta). Images were taken at different timepoints during infection and show the distribution of SLC7A5 and M2 in the cells. The maximum intensity projection was generated from z-stacks to visualise overall protein localisation. **g,** Pearson correlation coefficient between SLC7A5 and M2 in both the membrane and cytoplasm of HBEpCs cells. Correlation was calculated to assess the degree of colocalisation between these two proteins using Cell Profiler with 15 % threshold. Statistical analysis was performed using a Kruskal–Wallis test ($P < 0.0001$) followed by Dunn's multiple comparisons. Significant increases in colocalisation were observed at 12 and 24 hpi compared to mock (****$P < 0.0001$), while changes at 4 and 8 hpi were not significant. n = 31–68 cells per condition. **h,** Intensity profile of a representative plane showing the localisation of SLC7A5 and M2 in several HBEpCs cells at 12 and 24 hpi. The profile represents the fluorescence intensity across a cross-section of the cells.

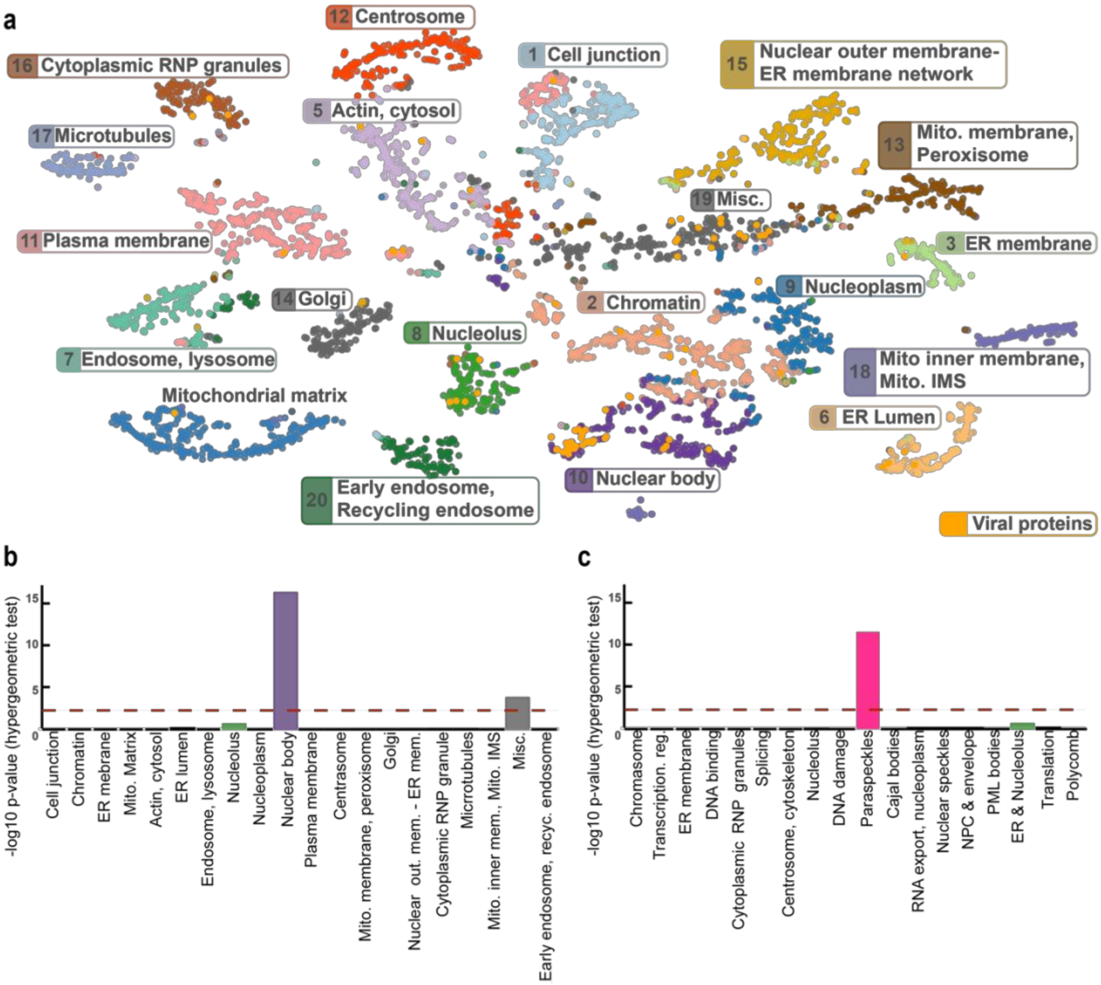


**Extended Data Fig. 8 | Spatial proteomic context of IAV–host protein interactions. a,** Mapping of host proteins crosslinked to IAV proteins onto the proximity-dependent biotinylation map of the human proteome[76]. Each coloured cluster represents a spatially resolved cellular compartment. Crosslinked host proteins were overlaid onto the map to infer their spatial context. **b,** Enrichment analysis of IAV-crosslinked host proteins using the Go et al. proximity labelling organelle map[76]. Significant enrichment was observed in the nuclear bodies (purple bar). **c,** Spatial enrichment using the updated nuclear body proteome dataset[77], revealing a strong enrichment in paraspeckles.

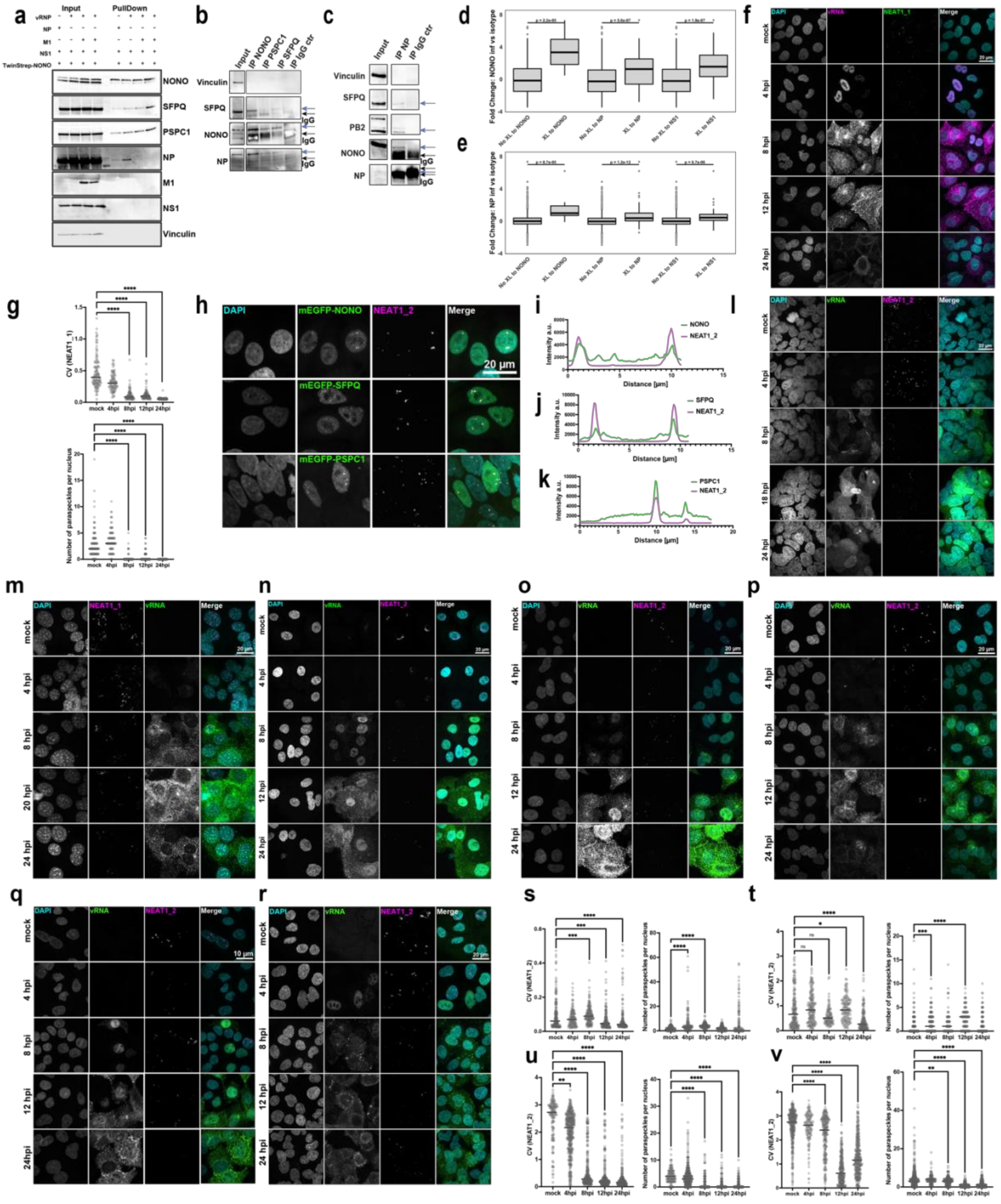


**Extended Data Fig. 9 | Analysis of paraspeckles during IAV infection. a,** TwinStrep-tagged NONO was overexpressed in HEK 293T cells alongside the viral proteins NP, M1, and NS1, or the vRNP complex for 24 hours. Streptactin beads were used for pull-down to isolate the TwinStrep-NONO complex. Input and pull-down samples were analysed by western blot. Antibodies against NONO, SFPQ, and PSPC1 were used to confirm successful pulldown of paraspeckle proteins. Antibodies against viral proteins NP, M1, and NS1 were employed to assess their interaction with the NONO complex. **b,** Immunoprecipitation of paraspeckle proteins NONO, SFPQ and PSPC1 and IgG control in A549 infected cells (WSN, MOI=3, 14 hpi). The proteins of interest are annotated with the blue arrows and the IgG bands with the black ones.

**Extended Data Video 1 | NONO or SFPQ cellular localisation in A549 cells stably overexpressing mEGFP-NONO or mEGFP-SFPQ infected with WSN at MOI 3 or mock-infected and imaged every 40 min**

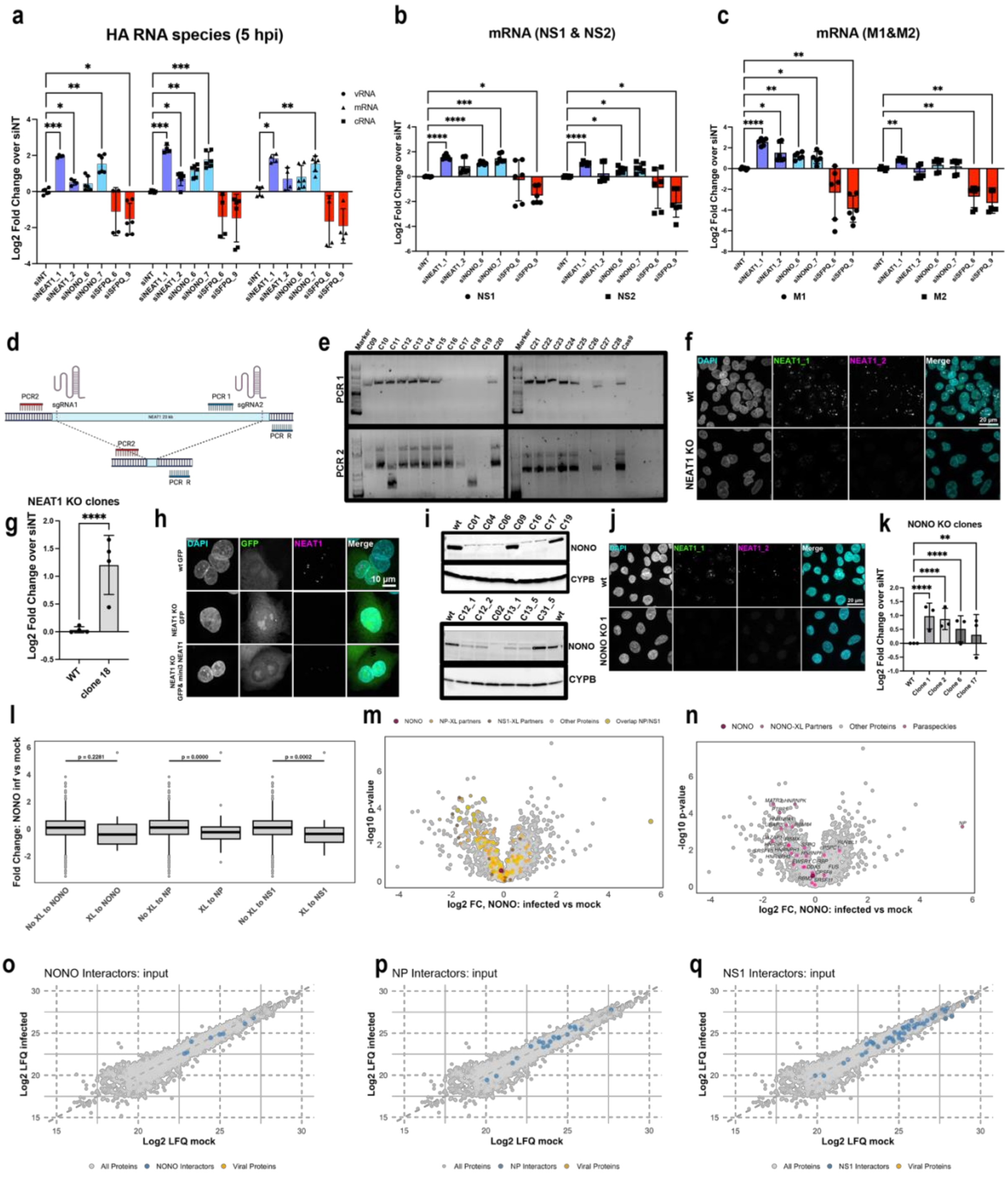


**Extended Data Fig. 10 | Role of NEAT1 and NONO during IAV infection. a,** qPCR analysis of viral RNA species of HA fragment normalised to GAPDH in A549 cells with knockdowns of paraspeckle proteins (WSN, MOI 3, 6hpi, n=3). **b-c,** qPCR analysis of spliced and unspliced viral RNA species in A549 cells with knockdowns of paraspeckle proteins (WSN, MOI 3, 6hpi, n=3): NS, NS2 (**b**) and M1, M2 (**c**). **d,** Strategy for NEAT1 KO Generation: Schematic of the CRISPR/Cas9-based approach to generate NEAT1 KO in A549 cells. Two sgRNAs are used to target NEAT1. PCR1 amplifies the region only in wild type (WT) or heterozygous clones, whereas PCR2 detects the shorter product (approximately 500 bp) in KO clones. For heterozygous clones, both the shorter KO product and the long (23 kb) WT product may appear. *Created in BioRender. https://BioRender.com/j78u936.* **e,** PCR Screening of Clones: Agarose gel showing PCR1 and PCR2 results for various clones. Clone 18 exhibits only the shorter PCR2 product, indicating a homozygous NEAT1 KO, while Clone 11 shows both products, indicating heterozygosity. **f,** Maximum projection of confocal images of NEAT1: Representative images showing the localisation of NEAT1_1 (green) and NEAT1_2 (magenta) in WT and NEAT1 KO (Clone 28) cells. DAPI (cyan) counterstain nuclei. Scale bar = 20 µm. **g,** Quantification of luciferase activity in WT and NEAT1 KO cells. **h,** Overexpression of miniNEAT1 3 in A549 NEAT1 KO cells using electroporation partially rescued paraspeckle formation. **I,** Western blot showing NONO protein levels in WT and NONO KO clones generated using sgRNA1 (above) sgRNA2 (below). CYPB serves as a loading control. **j**, Maximum projection of confocal images of NEAT1: IF images of NEAT1_1 (green) and NEAT1_2 (magenta) in WT and NONO KO (Clone 2) cells. DAPI (cyan) counterstains nuclei. Scale bar = 20 µm. **k,** Quantification of luciferase activity in WT and NONO KO A549 cells. **I,** Fold-change in NONO AP_MS (infected vs. mock) identified in AP-MS, including NONO, NP, and NS1 cross-linked partners. Data represent mean ± SD (n = 3). Statistical significance was determined using a Wilcoxon test. **m,** Volcano plot showing fold-change (log2 FC) vs. statistical significance (-log10 p-value) of NONO AP-MS in infected vs. mock samples. Paraspeckle proteins and NONO cross-linked partners are highlighted in pink. **n,** Volcano plot showing fold-change (log2 FC) vs. statistical significance (-log10 p-value) of NONO AP-MS in infected vs. mock samples. NP-XL, NS1-XL, and overlapping interactors are colour-coded. **o-q,** Scatter plots of Log2 LFQ values comparing infected and mock input samples in AP-MS (NONO inf and NONO mock). (o) NONO XL-partners, (p) NP XL-partners, and (q) NS1 XL-partners are highlighted in blue. Data from panels (**a-c,g,k**) represent mean ± SD (n = 3). Statistical analysis was performed using mixed effect model (a-c) the one-way ANNOVA (g,k) with Dunnett multiple comparison adjustment, two-sided. Significance levels: ***p < 0.001, **p < 0.01, *p < 0.05, ns = not significant.
